# D-serine suppresses one-carbon metabolism by competing with mitochondrial L-serine transport

**DOI:** 10.1101/2024.09.03.610855

**Authors:** Masataka Suzuki, Kenichiro Adachi, Pattama Wiriyasermukul, Mariko Fukumura, Ryota Tamura, Yoshinori Hirano, Yumi Aizawa, Tetsuya Miyamoto, Sakiko Taniguchi, Masahiro Toda, Hiroshi Homma, Kohsuke Kanekura, Kenji Yasuoka, Takanori Kanai, Masahiro Sugimoto, Shushi Nagamori, Masato Yasui, Jumpei Sasabe

## Abstract

L-serine serves as a central metabolic node that integrates glycolytic flux, lipid metabolism, and one-carbon metabolism. In the mature central nervous system, L-serine is actively stereo-converted to D-serine, which functions as a neurotransmitter. However, the role of D-serine in cellular metabolism remains unclear. Here, we show that D-serine competes with mitochondrial L-serine transport, thereby suppressing one-carbon metabolism. Metabolomic analysis revealed that D-serine reduces intracellular glycine and formate levels, indicating inhibition of the initial step of the one-carbon pathway. Molecular dynamics simulations and enzymatic assays revealed that D-serine has low affinity for serine hydroxymethyltransferase 2 (Shmt2), which catalyzes the first step in mitochondrial one-carbon metabolism, and does not directly inhibit its activity. Instead, membrane transport assays demonstrated that D-serine competes with mitochondrial L-serine transport, depleting the substrate of Shmt2. Functionally, under L-serine poor conditions in vitro and ex vivo, D-serine inhibited the proliferation of immature and undifferentiated neural cells including glioblastoma stem cells, which depend highly on one-carbon metabolism. Notably, endogenous D-serine levels were low during early neurodevelopment, but increased with maturation, coinciding with a shift in the transcriptional profiles of serine metabolic enzymes at the cellular level. Given that L-serine supports neurodevelopment and D-serine modulates neurotransmission, this developmental shift in serine enantiomer metabolism appears to align with the functional transitions of the maturing nervous system. Thus, our findings reveal that serine chirality can influence mitochondrial substrate availability and one-carbon flux, offering previously unappreciated insight into the stereoselective regulation of cellular metabolism.

## Introduction

Serine is a unique amino acid, both L- and D-enantiomers of which are synthesized *de novo* in the central nervous system. L-serine is primarily synthesized from a glycolytic intermediate through the phosphorylated pathway, and further stereo-converted to D-serine. Serine enantiomers and their metabolites are critically involved in neural development and neurophysiology in three major ways. First, L-serine is a crucial carbon donor with the cofactor, folate for the one-carbon metabolism, which is a universal metabolic process for nucleic acid synthesis, methylation, and reductive metabolism. As these pathways essentially support proliferative cells, undifferentiated or cancer cells are particularly susceptible to deprivation of L-serine or inhibition of *de novo* L-serine synthesis (Geeraerts et al., 2021). Second, biosynthesis of membrane lipids requires L-serine as an integral component of sphingolipids and phosphatidylserine. The nervous system includes highly polarized neurons and glia and exhibits the highest lipid abundance and complexity. Among membrane lipids, sphingolipids are particularly abundant in the nervous system and are essential for neuronal differentiation, polarization, synapse formation, and myelination, which are required for development and functional integrity of the nervous system(Hirabayashi and Furuya, 2008). Third, L-serine serves as a precursor for neurotransmitters, such as glycine and D-serine. L-serine is converted to glycine by serine hydroxymethyltransferase (Shmt) in the one-carbon metabolic pathway, and also into D-serine by serine racemase (Srr). Both glycine and D-serine bind *N*-methyl-D-aspartate (NMDA) receptor subunits with distinct affinities and regulate excitatory neurotransmission essential for neural development and function (Chatterton et al., 2002; Nancy and Dingledine, 1988). In addition, glycine also binds to inhibitory glycine receptors and controls early embryonic development as well as a variety of motor and sensory functions (Lynch, 2004).

Defects of L-serine synthesis due to single nucleotide polymorphisms in the genes encoding 3-phosphoglycerate dehydrogenase (PHGDH), the rate-limiting enzyme in the phosphorylated pathway, or the downstream enzyme phosphoserine aminotransferase (PSAT1) are associated with congenital microcephaly and hypomyelination, and exhibit psychomotor retardation and seizures in humans (Jaeken et al., 1997; KONING et al., 2002). Phgdh deficient mice die in embryo after 13.5 days post-coitum and show systemic growth defects, especially severe hypoplasia of the central nervous system, with marked reductions of L-serine and sphingolipids (Yoshida et al., 2004). Indeed, proliferative-marker-positive cells and mature neurons are almost entirely absent in these mice, indicating that *de novo* biosynthesis of L-serine is critical for neuronal proliferation and differentiation. Given its contribution to one-carbon metabolism, lipid synthesis, and production of glycine and D-serine (Yang et al., 2010), L-serine likely serves multiple functions in neural development. Of note, serine synthesis shows drastic changes after neurogenesis. Neural progenitor cells (NPCs) express PHGDH, whereas mature neurons lack its expression and lose the ability to synthesize L-serine. In mature cerebral cortex and hippocampus, on the other hand, astrocytes take over L-serine synthesis, and supply neurons with L-serine for synthesis of neurotransmitter D-serine (Wolosker, 2011). This metabolic compartmentalization suggests distinct functions of D- and L-serine across neural cell types and developmental stages. However, how D-serine contributes to cellular metabolism beyond neurotransmission has not been fully elucidated.

In this study, we used a metabolomics-based approach to examine how D-serine influences cellular metabolism, identified its primary target pathway in vitro, and assessed its functional consequences in neural cells both *in vitro* and *ex vivo*.

## Main

### D-serine impacts metabolites in one-carbon metabolism

To test whether serine chirality influences cellular metabolism in cortical neurons, we compared metabolomic profiles of primary cultured neurons (PCNs) treated with or without L- or D-serine for 48 h starting at DIV1. Serine enantiomers markedly altered the levels of metabolites such as glycine, cysteine, hypotaurine, taurine, glutathione, AMP, uracil, and polyamines (Figure 1A; Figure 1 –figure supplement 1A and B). These metabolites are components of the one-carbon metabolic network, including the folate cycle and synthetic pathways for purine, glutathione, taurine, and polyamine (Figure 1B), all of which are initiated by one-carbon donation from L-serine. Compared to D-serine, L-serine affected a broader range of metabolic pathways, including several amino acid metabolism (Figure 1C; Figure 1 –figure supplement 1C). Notably, however, D-serine uniquely caused a marked reduction in glycine levels (Figure 1A), which was further confirmed using a two-dimensional HPLC system (2D HPLC), a highly sensitive tool to quantify enantiomers, to occur independently of any change in L-serine levels (Figure 1E and F). Metabolites that showed similar reduction patterns to glycine were primarily polyamines and their intermediates (Figure 1D). Given that glycine is synthesized in the initial step of the one-carbon pathway, and that polyamines are produced in a downstream branch of the one-carbon network (Figure 1B), these results suggest that D-serine broadly suppresses one-carbon metabolism. In support of our view, D-serine treatment on PCNs further reduced production of formate, an intermediate of the folate cycle in one-carbon metabolism, whereas L-serine enhanced it (Figure 1G). Therefore, these findings indicate that D-serine exerts an inhibitory effect on the one-carbon metabolism, in contrast to the supportive role of L-serine.

**Figure 1.**
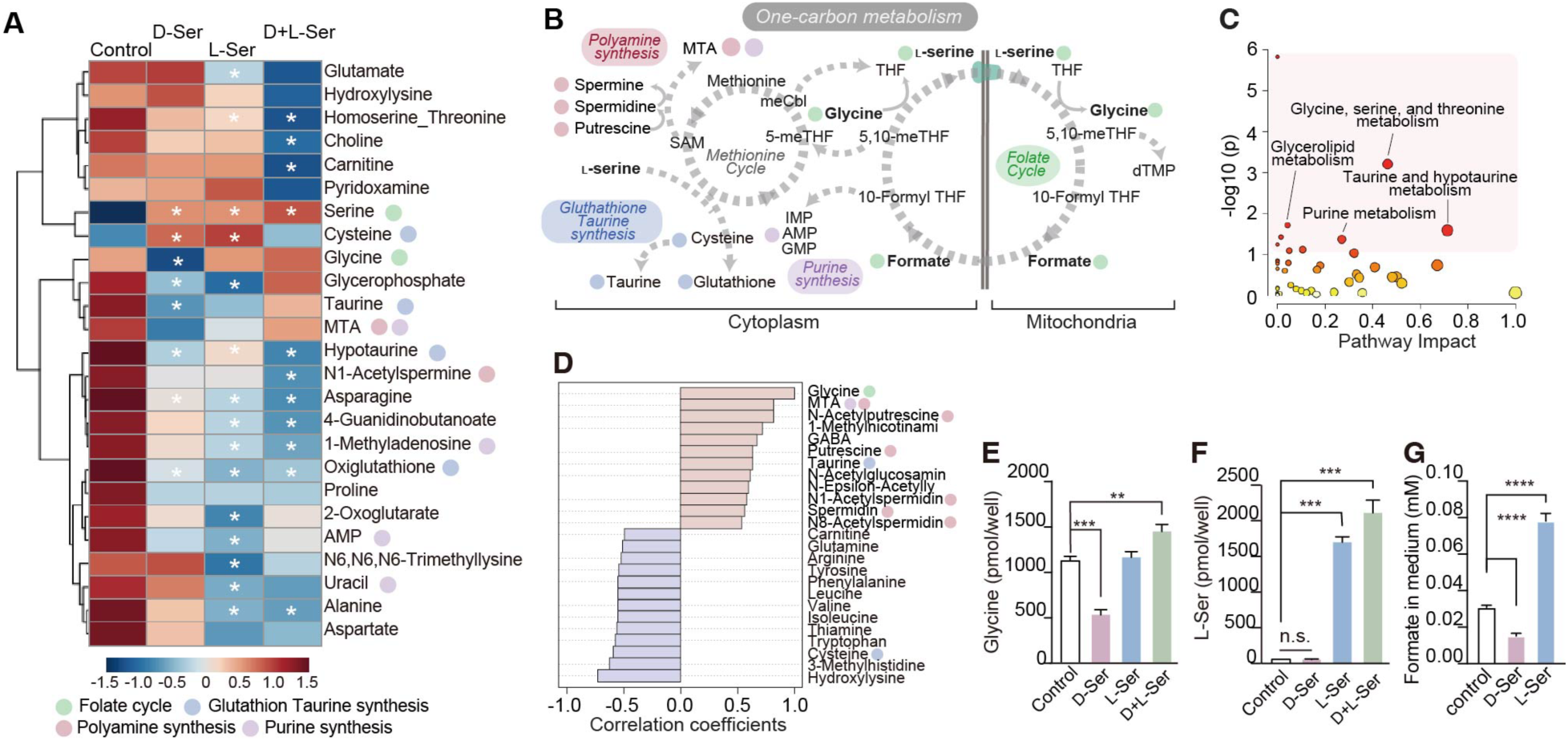
D-serine inhibits one-carbon metabolism. **A.** A heatmap shows top 25 metabolites in PCNs altered by treatment with D- and/or L-serine in the metabolomic analysis (n = 3). Color code represents z-score of the relative amount among conditions. Metabolites related to folate cycle (green), glutathione taurine synthesis (blue), polyamine synthesis (pink), and purine synthesis (purple) are indicated with colored circles. \**p* < 0.05 One-way ANOVA, compared to control. **B**. Major metabolites and pathways in the one-carbon metabolism. **C.** KEGG metabolic pathways enriched in D-serine treated cells compared to vehicle-control treated cells. The Pathway impact score is displayed on the X-axis and the -log10 transformed p-value on the Y-axis. **D.** Pearson’s correlation coefficients between glycine and other metabolites. Top 25 metabolites positively (red) and negatively (blue) correlated with glycine are shown. The colored circles indicate the relevant pathways, as in Figure 1A. **E, F**. Concentrations of glycine (**E**) and L-serine (**F**) in PCNs treated with D- and/or L-serine for 48 hours, were quantified using HPLC (n = 4). **G.** Concentrations of formate in the culture media of PCNs treated with 2 mM D- or L-serine for 48 hours were measured using an enzymatic assay (n = 4). Data are presented as the mean ± s.e.m. (**E, F,** and **G**). Statistical significance was evaluated by one-way ANOVA followed by Dunnett’s post hoc-test (**E, F,** and **G**). \**p* < 0.05 (**A**). Data are the representative of two independent experiments.

### D-serine suppresses one-carbon flux by competing with L-serine transport to mitochondria

In the central nervous system, glycine is synthesized from L-serine by serine hydroxymethyltransferase (Shmt) 1 and 2 (Figure 2A), which performs the first transfer of a carbon unit from L-serine to folate in one-carbon metabolism (Pan et al., 2020). In the presence of mitochondrial Shmt2, transfer of a carbon unit from L-serine to tetrahydrofolate (THF) and glycine production depends primarily on mitochondrial enzyme, but not cytoplasmic Shmt1 (Figure 2A)(Ducker et al., 2016). Therefore, the molecular target of D-serine in its inhibition of the initial step of one-carbon metabolism is likely mitochondrial Shmt2 or supply of its substrates into mitochondria (Figure 2A). To study whether D-serine impacts catalytic activity of Shmt2, we compared homotetrameric assembly of human SHMT2 combined with either L- or D-serine in the presence of THF and pyridoxal phosphate (PLP) (Figure 2 –figure supplement 2). When L- or D-serine was placed in the binding pocket of Shmt2, the eliminating carbon in D-serine was more distant from phosphorus (P) and oxygen (O) in PLP by 1.50 Å (C-P) and 1.10 Å (C-O) than that in L-serine (Figure 2B; Figure 2 –figure supplement 2A and B), suggesting that D-serine is not a good substrate for SHMT2. To investigate the stability of serine enantiomers in the binding pocket, we further performed molecular dynamic (MD) simulation of the serine-bound SHMT2 complex systems. MD simulation indicated that binding of D-serine did not affect the overall structure of SHMT2 compared to that of L-serine (Figure 2 – figure supplement 2C). On the other hand, molecular interactions of D-serine with SHMT2 were unstable and D-serine could not stay in the serine-binding pocket of SHMT2 (Figure 2C and Supplementary movie 1), which is supported by a recent report that SHMT does not use D-serine as a substrate in the canonical hydroxymethyl-transferase reaction in one-carbon metabolism (*K*m of SHMT2 = 0.07 mM for L-serine and 8.5 mM for D-serine) (Miyamoto et al., 2024). In contrast, L-serine remained placed stably in the pocket (Figure 2C and Supplementary movie 2). Therefore, D-serine was unlikely to compete with L-serine binding to SHMT2. Indeed, an *in vitro* experiment using recombinant SHMT2 showed that D-serine did not affect conversion of L-serine to glycine (Figure 2D).

**Figure 2.**
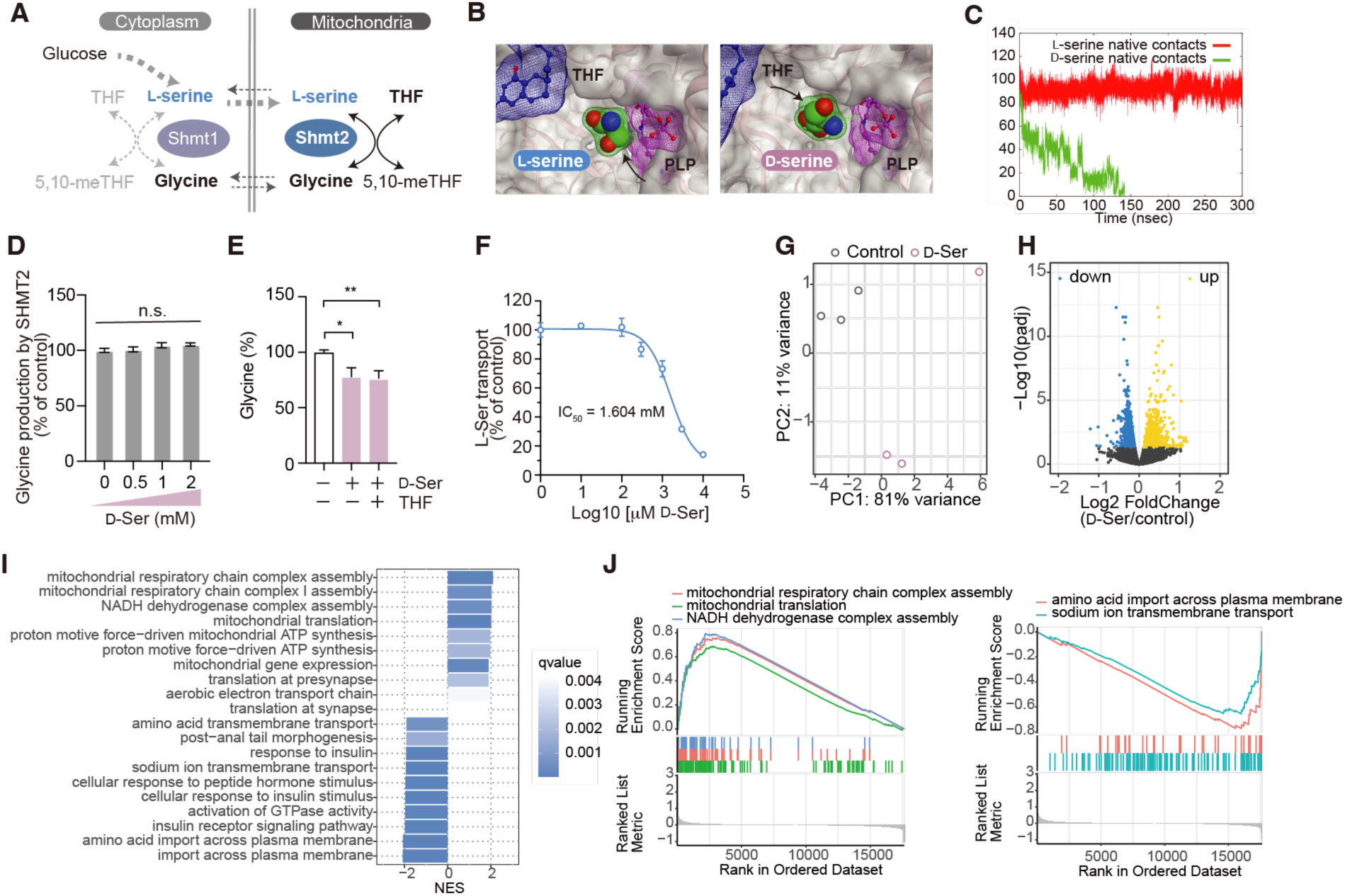
D-serine inhibits cytoplasmic L-serine transport to mitochondria. **A.** L-serine/glycine transport and conversion across cytoplasm and mitochondria in one-carbon metabolism. **B.** Binding sites of D-(right) or L-serine (left), THF, and pyridoxal phosphate (PLP) in the ligand-bound SHMT2 complex systems. Arrows indicate carbon atoms on serine enantiomers that transfer to THF. **C.** Time series variation of the number of contact atoms within the distance cutoff of 7.0 Å from D- or L-serine observed at the initial structure. **D.** SHMT2 enzymatic activity was determined. Glycine produced by purified SHMT2 from L-serine in the presence or absence of D-serine, was measured using HPLC. The result is shown as percent of control (with no D-serine) (n = 3). **E.** Relative concentrations of glycine in NPCs treated with or without 2mM D-serine and/or 50 μM THF were measured using HPLC. **F**. Inhibition of mitochondrial L-serine transport by D-serine was examined in semi-permeabilized cells (n = 4). The x-axis and y-axis represent the D-serine concentration used (log10-transformed), and the L-serine transport rate compared to the untreated control, respectively. **G.** A PCA plot shows RNA-seq results from cells treated with D-serine (pink) or control (gray). **H**. A volcano plot indicates differentially expressed genes in cells treated with D-serine vs controls. Gene expressions with adjusted *p*-values < 0.05 are highlighted in yellow (up) or blue (down). **I**. Bar plots show Normalized Enrichment Score (NES) for the top 10 up or down-regulated GSEA-enriched categories. The color code indicates the q-value. **J**. GSEA plots for a core subset of gene ontologies are displayed. The left panel shows pathways of mitochondrial respiratory chain complex assembly (red), mitochondrial translation (green), and NADH dehydrogenase complex assembly (blue). The right panel shows pathways of amino acid import across plasma membrane (red) and sodium ion transmembrane transport (cyan). Data are plotted as the mean ± s.e.m. (**D, E,** and **F**). Statistical significance was evaluated by one-way ANOVA followed by Tukey’s post hoc-test (**D**) or Dunnett’s post hoc-test (**E**).

Next, we wondered whether D-serine inhibits supply of Shmt2 substrate(s), L-serine or THF, to mitochondria. Since reduction of glycine caused by D-serine in PCNs was compensated by addition of L-serine, but not of THF (Figure 1E, 2E), we assumed that L-serine supply to mitochondria is disturbed by D-serine. Moreover, in PCNs, endogenous L-serine synthesis was not disturbed by addition of D-serine (Figure 1F), suggesting that D-serine affects L-serine transport to mitochondria, but not its synthesis. To confirm that D-serine impacts L-serine transport to mitochondria, we semi-permeabilized a neuroblastoma cell line, Neuro2a, to loosen cell membranes and monitored [H^3^]L-serine transport into mitochondria and other organelles in the presence or absence of D-serine. Intriguingly, D-serine competitively inhibited transport of L-serine in a dose-dependent manner (IC50, 1.604 mM) (Figure 2F). These results suggested that D-serine impairs one-carbon metabolism by competing with L-serine transport to mitochondria, but not by direct inhibition of Shmt2 activity. Sideroflexin 1 (Sfxn1), a multipass inner mitochondrial membrane protein, has been identified as a mitochondrial L-serine transporter (Kory et al., 2018) To assess whether Sfxn1 mediates the observed effect, we performed siRNA-mediated knockdown of Sfxn1. However, suppression of Sfxn1 alone did not significantly reduce L-serine transport, and D-serine retained its inhibitory effect under these conditions (Figure 2 –figure supplement 3). Given that Sfxn homologs (Sfxn1-Sfxn5) are proposed to function redundantly in mitochondrial serine transport (Kory *et al*, 2018), these results suggest that D-serine interferes with mitochondrial L-serine transport through multiple Sfxn family members rather than Sfxn1 alone.

To further understand the overall cellular responses to the D-serine burden, we performed RNA-sequencing using Neuro2a cells treated with D-serine, as these proliferative neural cells share metabolic characteristics with PCNs at early stages of differentiation. D-serine mildly, but significantly altered the transcriptome profile compared with vehicle control (Figure 2G and H). There were 685 up-regulated genes (log_2_ fold-change > 0, adjusted p-value < 0.05) and 905 down-regulated genes (log_2_-fold change < 0, adjusted p-value < 0.05) (Figure 2H). To determine whether genes with significant alterations are clustered in specific functional gene sets, we performed gene set enrichment analysis. Notably, D-serine enhanced expressions of genes associated with mitochondrial functions, such as respiratory chain complex assembly, ATP synthesis driven by proton motive force, and mitochondrial gene expression / translation. On the other hand, gene sets linked to amino acid import across the plasma membrane, cellular growth, and neuron projection extension were down-regulated (Figure 2I and J; Figure 2 –figure supplement 4). These dynamic transcriptional changes in the neuroblastoma cell line indicate that D-serine can impact polarization and growth of immature neurons negatively and further indicate that immature neurons resist D-serine-induced cellular stress by enhancing mitochondrial function, including energy synthesis.

### D-serine inhibits proliferation of tumor cells

One-carbon metabolism provides essential metabolites for nucleotide synthesis, methylation, and reductive metabolism, and supports cellular proliferation. Tumor cells are highly proliferative and are particularly susceptible to deprivation of one-carbon units by L-serine restriction or inhibition of *de novo* serine synthesis (Newman and Maddocks, 2017). Given that D-serine inhibits one-carbon flux by competing with L-serine in PCNs and the neuroblastoma cell line, we wondered whether D-serine influences proliferation of neural tumor cells. To test this idea, we treated several human/rodent neuroblastoma cell lines cultured in the regular medium containing sub-millimolar L-serine and fetal bovine serum with D-serine. Consistent with our findings that D-serine competes with L-serine transport (Figure 2F), high concentrations of D-serine were required to suppress proliferation of neuroblastoma cell lines cultured in regular medium (Figure 3 –figure supplement 5A). In contrast, culturing in L-serine/glycine-free medium with dialyzed fetal bovine serum reduced the proliferation of Neuro2a cells with much lower D-serine concentrations (IC50 = 1.94 mM) (Figure 3 –figure supplement 5B-D), which corresponds to the efficacy of D-serine on inhibition of L-serine transport (IC50 = 1.604 mM) (Figure 2F). Indeed, with L-serine/glycine-free medium, D-serine significantly inhibited proliferation of all neuroblastoma cell lines tested, at low millimolar concentrations (Figure 3A). In accordance with its effect on PCNs (Figure 1E), D-serine inhibited production of glycine and suppressed proliferation of Neuro2a cells, which was rescued by supplementation with L-serine (Figure 3B; Figure 3 –figure supplement 5E). Moreover, supplementations with glycine and downstream metabolites of the folate cycle in the one-carbon pathway (Figure 1B), such as formate, inosine monophosphate (IMP), or 5,10-meTHF, phenocopied the rescue effect by L-serine (Figure 3 –figure supplement 5E-I). On the other hand, combinations of glycine and metabolites of the methionine cycle (methionine or methylcobalamin (meCbl)) as well as other metabolites associated with one-carbon metabolism (thymidine monophosphate (dTMP) or glutathione) (Figure 1B) failed to liberate cells (Figure 3 –figure supplement 5J and K). These results suggest that D-serine impacts the folate cycle rather than the methionine cycle in one-carbon metabolism to prevent growth of neuroblastoma cells.

**Figure 3.**
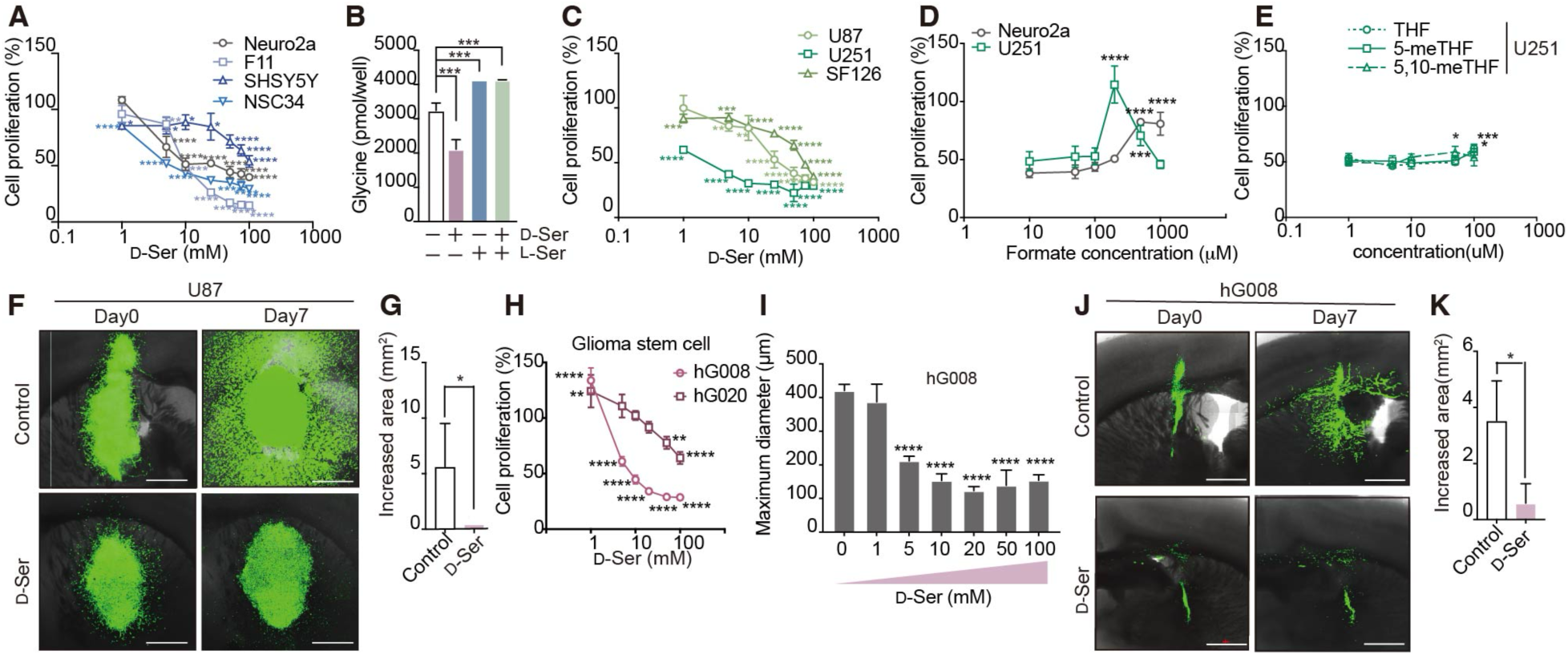
D-serine attenuates proliferation of neural tumor cells A, C, **H**. Relative proliferation of neuroblastoma cell lines (**A**), glioblastoma lines (**C**), and glioma stem cells (**H**) treated with D-serine at indicated doses were analyzed 48 h after treatment (n = 4, each). Cells with the vehicle treatment were used as 100% proliferation controls. **B**. Glycine concentration in Neuro2a cells was quantified by HPLC at 48 h after treatment with D- and/or L-serine (n = 3). **D, E**. Cell proliferation was assessed at 48 h after treatment with 5 mM D-serine and formate/folates at indicated doses (n = 4). Cell proliferation was compared to that of cells treated with D-serine alone. **F, G, J, K.** *Ex vivo* slice cultures of mouse brain transplanted with U87 (**F, G**) or hG008 cells (**J, K**) expressing Venus fluorescence were treated with D-serine. Representative images of the xenograft at 0 and 7 days after treatment are shown (**F, J**). Increased area of fluorescence after 7 days of culture was measured (**G, K**). **I**. Sphere diameters of glioma stem cells (hG008) were measured at 7 days after treatment with D-serine at indicated doses. Data are presented as the mean ± s.e.m. (**A-E, G-I,** and **K**). Statistical significance was evaluated by one-way ANOVA followed by Dunnett’s post-hoc test (**B, D,** and **E**) or by Student’s t-test (**G** and **K**).

As neurons and astroglia share a common developmental origin, we further tested whether glial tumor cells also show vulnerability to D-serine. Glioblastoma is the most common malignant brain tumor in adults, whose survival rate is low despite advances in surgical and medical neuro-oncology(Bent et al., 2023; Delgado-López and Corrales-García, 2016). Among human glioblastoma cell lines (U87, U251, SF126), U251 cells were the most sensitive to D-serine treatment (Figure 3C). Supplementation with glycine and formate restored proliferation of U251 (Figure 3D), and 5,10-meTHF recovered the vulnerability to D-serine, albeit very mildly (Figure 3E), supporting the significance of one-carbon metabolism in growth of glioblastoma cell lines too. To further test the anti-proliferative effect of D-serine in the tumor microenvironment, we made an organotypic brain slice culture from mice with orthotopic xenografts of glioblastoma cells expressing Venus fluorescence(Tamura et al., 2019). Brain slices of mice implanted with U87 cells in the striatum were cultured *ex vivo* for 7 days in serine/glycine-free media containing D-serine or vehicle. D-serine strikingly suppressed expansion of U87 cells with Venus fluorescence compared to the control treatment, even in *ex vivo* tumor culture (Figure 3F and G). Since glioma stem cells (GSCs) in the glioblastoma have self-renewal and tumor-initiating capacity and cause cancer recurrence(Gimple et al., 2022; Tang et al., 2021), we wondered whether GSCs also have similar vulnerability to D-serine. *In vitro*, D-serine reduced viability of cell lines for GSCs (hG008 and hG020), established from human glioblastoma specimens^25^, in a dose-dependent manner with greater efficacy to the hG008 line (Figure 3H). This growth defect in hG008 cells induced by D-serine was rescued by supplementation with glycine and formate, as in neuroblastoma and glioblastoma cells (Figure 3 –figure supplement 5L). hG008 cells have a sphere-forming capacity^23^, and we also observed D-serine-induced reduction of sphere diameters of hG008 cells cultured under the sphere-forming conditions (Figure 3I). Moreover, in brain slices of mice with orthotopic xenografts of hG008 cells expressing Venus fluorescence, D-serine significantly inhibited growth and invasiveness of hG008 cells (Figure 3J and K). Thus, brain tumor cells showed consistent vulnerability to D-serine via inhibition of one-carbon metabolism.

### D-serine induces apoptosis by interfering with one-carbon metabolism in neuroprogenitor cells

Neural stem cells have similar characteristics to neural tumor cells with high proliferative capacity(Reya et al., 2001). To test whether D-serine also affects immature neurons, we made a primary culture of neuroprogenitor cells (NPCs) from telencephalon in E14 mice in serine-free Neurobasal medium, treated with D- or L-serine from DIV1. At DIV7, L-serine treatment significantly increased the number of neurons with a condensed neurites compared to vehicle-treated neurons. In contrast, D-serine-treated culture had fewer number of neurons with sparse neurites, suggesting that D-serine has a negative impact on maturation of NPCs (Figure 4A). Furthermore, D-serine activated cleavage of caspase-3 dose-dependently in Nestin-positive NPCs and Tubb3+ developing neurons 48 h after treatment, whereas L-serine had no effect (Figure 4B-D; Figure 4 –figure supplement 6A-C –Source Data 1). In contrast, vesicular glutamate transporter 2 (vGlut2)-positive mature excitatory neurons, glutamate decarboxylase-67 (Gad67)-positive mature inhibitory neurons, and glial fibrillary acidic protein (Gfap)-positive glial cells did not undergo apoptosis under D- or L-serine treatment (Figure 4B; Figure 4 –figure supplement 6D and E), suggesting that mature neurons and astrocytes are resistant to D-serine. Consistently, D-serine treatment at DIV7 did not activate cleavage of caspase-3 or reduce proliferation of neurons (Figure 4E and F–Source Data 1). Susceptibility of NPCs was pronounced in the presence of D-serine, but was not caused by other D-amino acids, such as D-aspartate, D-alanine, or D-proline (Figure 4 –figure supplement 7A –Source Data 2), which can be detected in mammalian intestinal bacteria(Gonda et al., 2023). As D-serine is a coagonist of NMDA receptors(Mothet et al., 2000; Wolosker and Balu, 2020), we further tested whether D-serine neurotoxicity involves NMDA receptors(McNamara and Dingledine, 1990). Blockade of Ca^2+^ influx through NMDA receptors by a non-competitive NMDA antagonist, MK-801, or of D-serine-binding to the receptors by 5,7-dichlorokynurenic acid (DCKA) did not attenuate D-serine neurotoxicity (Figure 4 –figure supplement 7B and C –Source Data 2). Also, inhibition of the receptor’s downstream signal by *N*-nitro-L-arginine methylester (L-NAME) or an antioxidant (glutathione) did not affect enhanced cleavage of caspase-3 by D-serine (Figure 4 –figure supplement 7D and E –Source Data 2), although D-serine decreased cellular glutathione (Figure 4 –figure supplement 7F). Thus, our findings suggest that apoptosis triggered by D-serine in NPCs does not mediate NMDA receptors. To test whether apoptosis triggered by D-serine involves amino acid metabolism, we supplemented NPCs treated with D-serine from DIV1 with various L-amino acids or glycine. As expected, among L-amino acids and glycine, L-serine specifically and dose-dependently inhibited cleavage of caspase-3 caused by D-serine (Figure 4G and H –Source Data 1), which is consistent with competitive mitochondrial transport of serine enantiomers (Figure 2F). Moreover, similar to the observation that 5,10-methylene THF rescued neural tumor cell growth (Figure 3 –figure supplement 5I), 5,10-methylene THF restored D-serine-induced cleavage of caspase-3 in NPCs (Figure 4I –Source Data 1), suggesting that D-serine interferes with L-serine functions, such as its role in one-carbon metabolism.

**Figure 4.**
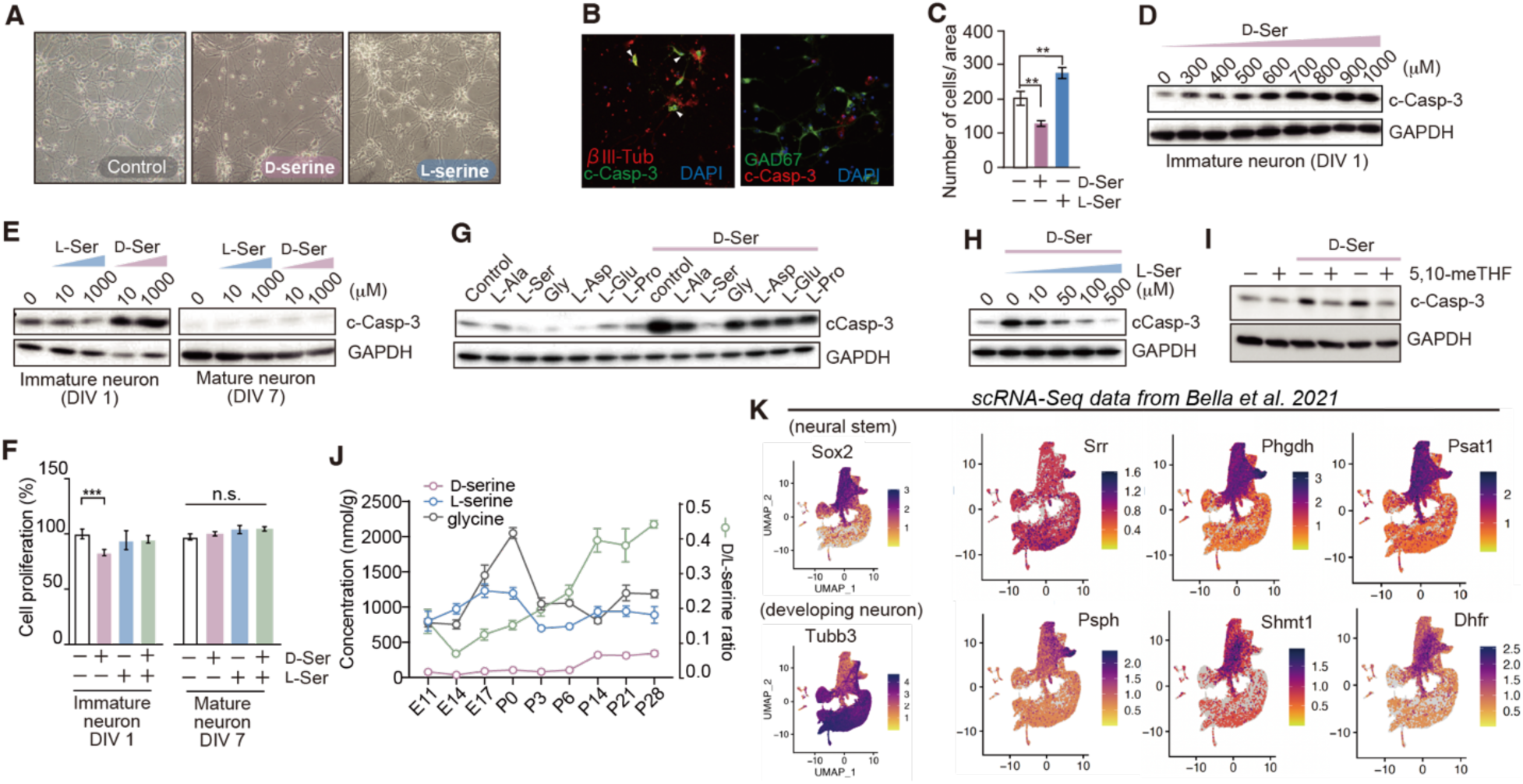
Developmental transition of serine enantiomer synthesis and enantio-selective effects on immature and mature neurons. **A**. Light microscopic view showing primary cultured NPCs treated with 1mM D- or L-serine for 7 days in serine-free media. **B.** Immunofluorescent images are NPCs treated with 1 mM D-serine for 48 hours. The left panel shows the co-staining with antibodies against cleaved caspase-3 (c-Casp-3, green), βIII Tubulin (red), and DAPI (blue). The right panel shows the co-staining with antibodies against c-Casp-3 (red), GAD67 (green), and DAPI (blue). **C**. Numbers of NPCs per area at DIV7 in (**A**) were counted (n = 4). **D.** Western blots indicate cleavage of caspase-3 in cultured neurons at 48 h after treatment of D-serine with indicated doses. **E.** Western blots indicate cleavage of caspase-3 in cultured neurons at 48 h after treatment of serine enantiomers with indicated doses starting at DIV1 (immature) or DIV7 (mature neurons). **F.** Relative cell proliferation of NPCs after 48 h treatment of 1mM serine enantiomers was analyzed (n = 4). **G. H.** Cleavage of caspase-3 in NPCs induced by 1mM D-serine was observed in the presence of various L-amino acids (1 mM each) (**G**) or L-serine at indicated doses (**H**). **I**. Cleavage of caspase-3 in NPCs triggered by 2 mM D-serine was observed in the presence or absence of 40 μM 5,10-meTHF. **J**. Concentrations of serine enantiomers and glycine and the ratio of serine enantiomer concentrations in the cerebrum during development (n= 3-4). **K**. Single-cell transcriptome profiles of enzymes involved in serine metabolic pathways during mouse brain development. Original data were from Bella et al (Bella et al., 2021). Data are plotted as the mean ± s.e.m. Statistical significance was evaluated by One-way ANOVA followed by Dunnett’s post hoc test (**C**). All *in vitro* data are the representative of at least two independent experiments.

To further rule out alternative mechanisms, we examined whether D-serine disrupts L-serine-mediated lipid synthesis. During the neuronal development, NPCs require L-serine to synthesize membrane sphingolipids and phospholipids that support cellular expansion and neurite elongation (Figure 4 –figure supplement 8A). A comparative lipidomic study using NPCs revealed that D-serine treatment increased phosphatidylserine, but not sphingolipids or other phospholipids (Figure 4 –figure supplement 8B and C). To test whether D-serine is incorporated into phosphatidylserine, we extracted membrane lipids, hydrolyzed phosphatidylserine using phospholipase D (PLD) to release serine, and quantified serine enantiomers using 2D HPLC (Figure 4 –figure supplement 8D and E). Phosphatidylserine extracted from NPCs cultured without D-serine harbored only L-serine, but supplementation with D-serine significantly increased phosphatidyl-D-serine and decreased phosphatidyl-L-serine (Figure 4 –figure supplement 8F and G). This effect was abolished by co-supplementation with L-serine (Figure 4 –figure supplement 8F and G). However, exogenous phosphatidyl-L-serine failed to rescue D-serine-induced apoptosis (Figure 4 –figure supplement 8H –Source Data 3), indicating that composition of membrane phosphatidyl-serine is not the primary cause of cell death. Together, these results demonstrate that D-serine impairs the survival of immature neural cells not by modulating neurotransmission or lipid metabolism, but by disrupting one-carbon metabolism through competition with L-serine.

### Enantiomeric shift of serine metabolism during neural development

Serine metabolism is fundamental to multiple cellular functions and crucial for neural development. To gain a functional overview of how serine chirality contributes to these processes, we monitored serine enantiomers and glycine using the 2D HPLC system. L-serine levels in the mouse cerebral cortex showed a gradual increase during embryonic development, dropped at birth, and remained at a concentration above 500 nmol/g thereafter (Figure 4J). As glycine synthesis depends in part on L-serine, glycine showed a similar trend to L-serine in embryo and maintained levels above 800 nmol/g after birth (Figure 4J). On the other hand, D-serine was trace during the embryonic stage, but increased gradually after birth. Therefore, the D/L-serine ratio exhibited a steady increase after birth and reached 0.4 in mature mouse brain (Figure 4J). The dynamics of serine enantiomers during the brain development is corroborated by transcriptional profiles associated with serine chial metabolism. Bulk RNA-seq data of developing neurons derived from mouse embryonic stem cells (ESC)(Hubbard et al., 2013) or human inducible pluripotent stem cells (iPSC)(Burke et al., 2020) in public databases showed that diverse transcriptions found in the phosphorylated pathway of L-serine synthesis (PHGDH, PSAT1, and PSPH), one-carbon metabolism (SHMT1/2, MTHFD1/2, and MTHFD1/2L), and nucleic acid synthetic pathway (DHFR, TYMS, MTHFS) decrease along with the neuronal maturation (Figure 4 –figure supplement 9A and B). In contrast, D-serine-synthetic enzyme SRR increases after days *in vitro* 7 (DIV7) in ESC-derived neurons and NPC/Rosetta cells (DIV14-21) in human iPSC-derived neurons. Subunits of NMDA receptors, including Grin1/GluN1, to which D-serine or glycine binds, are expressed in matured neurons after DIV16 of mouse ESC-derived cells (Figure 4 –figure supplement 9A) or those at DIV49 of human iPSC-derived cells (Figure 4 –figure supplement 9B). Alterations of serine metabolism during development were further supported by single-cell RNA-seq performed by Bella et al.(Bella et al., 2021), which provides crucial information about the trajectory of differentiating cells and cell type-specific transcriptional profiles during mouse brain development *in vivo*. Uniform manifold approximation and projection (UMAP) illustrates that Sox2+ neural progenitor cells (cluster 1, 7, 9, and 11) express L-serine synthetic enzymes (Phgdh, Psat1, Psph) and folate cycle enzymes (Tyms, Mthfd1, Dhfr) as well as a proliferation marker gene Mki67 (Figure 4K; Figure 4 –figure supplement 9C and D). Apoe+ and Aldh1l1+ astrocytes (cluster 7) also express L-serine synthetic pathways, whereas Tubb3+ mature neurons (cluster 0, 2, 3, 4, 5, 8, 12, 13, 15, 16, 19, 22) do not express serine synthetic enzymes (Figure 4K; Figure 4 –figure supplement 9C and D). Tubb3+ neurons weakly express Srr at P4 but not E10 (Figure 4 –figure supplement 9E). These transcriptional profiles suggest that reduction of L-serine biosynthesis during neural development accompanies inactivation of the folate cycle and one-carbon metabolism, and that mature neurons start to synthesize D-serine. Given that the folate cycle and one-carbon metabolism, essential for nucleic acid biosynthesis, are the metabolic signature of cell proliferation, NPCs lose their metabolic activity for proliferation in the embryonic stage but instead develop the capacity for neurotransmission during the postnatal development. Although we found that D-serine inhibits L-serine-dependent one-carbon metabolism, its endogenous production is minimal in neural stem/progenitor cells and increases only after the onset of neuronal maturation, coinciding with the upregulation of Srr. Thus, D-serine is unlikely to play a direct role in regulating proliferative metabolism during early neurodevelopment. Rather, its selective synthesis in post mitotic neurons appears metabolically rational, aligning with their shift from proliferation to neurotransmission.

## Discussion

We found that D-serine competes with mitochondrial transport of L-serine, thereby limiting substrate availability for Shmt2 and suppressing one-carbon metabolic flux (Figure 4 –figure supplement 10). This metabolic interference reduces glycine and formate levels and alters downstream pathways including polyamine synthesis. Functionally, under L-serine limited conditions, D-serine suppresses the proliferation and survival of immature neural cells, independent of NMDA receptor activation. Moreover, we demonstrated that endogenous D-serine production increases postnatally in tandem with neuronal maturation, whereas early progenitor cells remain devoid of D-serine synthetic enzyme. In addition to the known role of D-serine in the functional maturation of neurons, our findings identify a previously unrecognized stereoselective role of D-serine in cellular metabolism, and provide a rationale for its selective synthesis in mature neurons where proliferative metabolic activity is no longer required.

In the central nervous system, L-serine increases during fetal development and declines rapidly after birth, while D-serine increases only after birth and reaches the mature levels within a few weeks (Figure 4J). This postnatal increase of D-serine is consistent with a previous report in rats (Hashimoto et al., 1993) and can be explained by a gradual increase of Srr after birth in mouse glutamatergic neurons (Figure 4 – figure supplement 9A and E) or forebrain tissue (Folorunso et al., 2021; Miya et al., 2008; Wang and Zhu, 2003). In the mature brain, the majority of Srr is restricted to neurons (Balu et al., 2014; Miya et al., 2008), which is consistent with the idea that D-serine synthesis accompanies neuronal maturation. In contrast, PHGDH is not expressed in neurons, but is confined to astrocytes in mature brain (Yang et al., 2010). This astrocyte-specific expression of the L-serine synthetic enzyme accords with our findings that a set of genes encoding enzymes for L-serine synthesis and one-carbon metabolism are downregulated in mature neurons, but confined to a cell population with mature astrocyte markers (Figure 4K; Figure 4 –figure supplement 9). Given that astrocytes require one-carbon metabolism for proliferation and that neurons prioritize neurotransmission, such cell-type specific and enantio-selective serine metabolism in the matured brain appears to be a reasonable metabolic compartmentalization. Indeed, loss of D-serine synthesis does not trigger developmental abnormalities (Miya et al., 2008), but causes functional or structural abnormalities in mature neurons (Balu et al., 2012; Basu et al., 2009; Horn et al., 2017; Sultan et al., 2015). Furthermore, supplementation with D-serine in the neonatal brain enhances functional development of forebrain neurons (Nomura et al., 2016). These findings suggest that D-serine primarily contributes to functional maturation rather than differentiation of NPCs.

While D-serine is important for functional maturation of neurons as well as excitatory neurotransmission through NMDA receptors(Basu et al., 2009; Horn et al., 2017; Mothet et al., 2000), D-serine has an inhibitory effect on proliferation and triggers apoptosis in NPCs and other immature cells (Figure 3 and 4). Extracellular D-serine is a classic trigger of excitotoxicity in neurons (Patel et al., 1990; Shleper et al., 2005), whereas it induces intracellularly necrosis/apoptosis in renal tubular cells at high doses (Kaltenbach et al., 1979; Williams and Lock, 2004). Excitotoxicity of D-serine has been reported in mature neurons in ischemic or neurodegenerative diseases and mediates NMDA receptors (Inoue et al., 2008; Mustafa et al., 2010; Sasabe et al., 2007; Shleper et al., 2005). In contrast, the D-serine toxicity, which we observed in neural culture, occurs only in immature neurons, and can be rescued by L-serine or intermediates of the one-carbon metabolism, but not by inhibitors of NMDA receptors or their downstream signals (Figure 4G-I and S7). The latter toxicity reported in kidneys mediates hydrogen peroxide generated through degradation of D-serine by a D-serine catabolic enzyme, D-amino acid oxidase, plentiful in kidneys (Maekawa et al., 2005). Nonetheless, D-amino acid oxidase is not expressed in neurons or in the forebrain (Gonda et al., 2022). Furthermore, as we could not find protective effects by an antioxidant against D-serine toxicity in immature neurons (Figure 4 –figure supplement 7E), involvement of oxidative stress by hydrogen peroxide is unlikely. Thus, D-serine toxicity observed here appears mechanistically distinct from previously reported forms. Importantly, Okada et al., suggests that renal tubular toxicity by D-serine can be ameliorated by L-serine (Okada et al., 2017). Although the authors did not demonstrate involvement of one-carbon metabolism in D-serine toxicity in the kidneys, their observation could be associated with our findings that D-serine functionally competes with L-serine (Figure 4H).

L-serine can fuel one-carbon metabolism through both cytoplasmic Shmt1 and mitochondrial Shmt2; however, the relative contribution of these pathways is likely to depend on cell type, differentiation state, and proliferative demand. Consistent with this notion, the differential sensitivity of immature and mature primary cortical neurons to D-serine (Figure 4E) may reflect differences in compartmentalized one-carbon metabolism. On the other hand, we demonstrated that D-serine does not directly inhibit Shmt2 activity (Figure 2D). Given that both Shmt1 and Shmt2 exhibit strong stereoselectivity for L-serine and do not utilize D-serine as a substrate (Miyamoto et al., 2024), the suppression of one-carbon metabolism observed in this study is more likely explained by altered mitochondrial L-serine transport rather than direct inhibition of Shmt reactions. Future isotope tracing studies using labeled L-serine will be valuable to dissect how D-serine influences the balance between cytosolic and mitochondrial one-carbon fluxes. Notably, the mitochondrial branch of one-carbon metabolism plays a critical role during embryonic development (Ducker and Rabinowitz, 2017; MacFarlane et al., 2008), highlighting mitochondrial L-serine transport as a key regulatory step for cellular proliferation during development. A seminal study by Kory et al. demonstrated that SFXN1 and its homologs mediate L-serine transport to mitochondria(Kory et al., 2018), and further showed *in vitro* that D-serine competes with L-serine transport through SFXN1. Together with our *in vivo* L-serine competition assays performed in the presence and absence of Sfxn1 (Figure 2F and S3), these findings support a model in which D-serine suppresses mitochondrial one-carbon metabolism by competing with mitochondrial L-serine transport mediated by multiple members of the Sfxn family rather than Sfxn1 alone.

We show that D-serine attenuates cell proliferation by inhibiting one-carbon metabolism (Figure 3). The concentration of D-serine required to inhibit proliferation in our system exceeds reported physiological concentration (∼300 μM). However, local intracellular concentrations of D-serine may be substantially higher than those estimated from homogenized cerebral tissue, particularly because Srr is localized within neuronal cytoplasm (Miya et al, 2008). Moreover, elevated D-serine levels have been reported in adult-onset neurodegenerative disorders including amyotrophic lateral sclerosis and Alzheimer’s disease, in which D-serine degradation is impaired (Madeira et al., 2015; Mitchell et al., 2010; Sasabe et al., 2012, 2007). Adult neurogenesis is reduced in these pathological conditions (Cao et al., 2024; Galán et al., 2017; Moreno-Jiménez et al., 2019), raising the possibility that dysregulated D-serine metabolism contributes to impaired neurogenesis (Sultan et al., 2013; Zhao et al., 2019). Nevertheless, proliferating neural progenitor cells express little Srr, making it unlikely that physiological D-serine concentrations alone determine cell fate during normal development. Rather, our findings suggest that the progressive increase of D-serine during brain maturation may contribute to a developmental metabolic shift that interferes with efficient L-serine-dependent one-carbon metabolism in postmitotic neurons. One-carbon metabolism is essential for nucleotide synthesis and methylation in proliferative tissues(Ducker and Rabinowitz, 2017), and immature neural cells appear particularly susceptible to D-serine-mediated metabolic perturbation (Figure 4). Consistent with the importance of this pathway during development, deficiencies of either L-serine or folate lead to congenital abnormalities of the central nervous system in humans(Acuna-Hidalgo et al., 2014; Beaudin and Stover, 2009). Furthermore, *de novo* serine synthesis and mitochondrial one-carbon pathway are required for the proliferative phenotype in a large variety of human cancers (Jain et al., 2012; Labuschagne et al., 2014; Lee et al., 2014; Lewis et al., 2014; Nilsson et al., 2014). The broad anti-proliferative effects of D-serine on neural tumor cells (Figure 3) are consistent with the conclusion that D-serine suppresses mitochondrial one-carbon metabolism. Although we tested only neural tumor models, the underlying mechanism may extend to wide range of cancers with varying dependence on mitochondrial one-carbon metabolism. Since cancer genetics indicate two pathways not targeted by existing therapies, de novo serine synthesis and mitochondrial one-carbon metabolism (Ducker and Rabinowitz, 2017; Newman and Maddocks, 2017), inhibition of mitochondrial L-serine transport to disconnect these two pathways may represent a potential therapeutic strategy. However, our exploratory *in vivo* experiments showed that high-dose D-serine administration caused behavioral abnormalities, suggesting that direct administration of D-serine itself may have limited therapeutic applicability. Future studies may therefore explore optimized D-serine derivatives or delivery strategies that preserve metabolic effects while reducing NMDA receptor-mediated activity.

The role of D-amino acids in cellular metabolism and neural development is only beginning to be understood. In addition to D-serine, Srr is capable of producing other D-amino acids, including D-cysteine, raising the possibility that multiple enantiomers contribute to developmental and metabolic regulation. Recent studies suggest both overlapping and distinct functions of D-serine and D-cysteine. For example, neural deletion of Srr reduces adult neurogenesis and alters lipid metabolism in the subventricular zone (Roychaudhuri et al., 2023) whereas embryonic Srr knockout increases Sox2^+^ neural progenitor cells, a phenotype proposed to involve D-cysteine rather than D-serine. Consistent with this idea, D-cysteine levels peak during early embryogenesis and decline during development, in contrast to the postnatal increase in D-serine (Semenza et al., 2021). Beyond neural development, D-cysteine has also been implicated in broader metabolic regulation, including tumor suppression through inhibition of cysteine desulfurase (NFS1) (Zangari et al., 2025) and modulation of insulin secretion via DNA methylation (Roychaudhuri et al., 2024). Given the close connection between L-serine metabolism, transsulfuration pathways, and S-adenosyl-methionine (SAM)-dependent methylation reactions, D-serine-mediated inhibition of mitochondrial L-serine transport may also influence cysteine and methylation metabolism. However, supplementation with SAM, methionine, or glutathione did not rescue D-serine-induced growth inhibition in our system (Figure 3 –figure supplement 5J), suggesting that these pathways are not the primary drivers of the observed phenotype. Together, these findings suggest that developmental-stage-dependent production of D-serine and D-cysteine by Srr may contribute to coordination of proliferation, differentiation, and metabolic state during neural development and disease.

Neuronal metabolism exhibits the unique property of serine isomerization to adapt to functional maturation (Figure 4). Consequently, mature cerebrum contains sub-millimolar levels of D-serine, which account for a quarter of total serine in the cerebrum and are about 100-times higher than blood levels(Miyoshi et al., 2009). This active stereo-conversion of serine in the cerebrum is unique to mammals among vertebrates(Nagata et al., 1994), implying that neurophysiological functions of D-serine are inseparable from the functional evolution of the cerebrum in higher vertebrates. Indeed, whereas either D-serine or glycine are required to activate NMDA receptors, D-serine binds to the GluN1 subunit of NMDA receptors with higher affinity than does glycine(Furukawa and Gouaux, 2003) and also has an inhibitory effect on the GluN3 subunit by competing with glycine(Chatterton et al., 2002). Therefore, extracellular D-serine appears to be evolutionarily important as a neurotransmitter, with different properties than glycine. On the other hand, the inhibitory function of intracellular D-serine against mitochondrial one-carbon metabolism (Figure 1 and 2) may be an evolutionarily inherited trait from the biological ancestors of mitochondria, bacteria(Archibald, 2015). Notably, bacteria depend on one-carbon metabolism for their growth, and D-serine attenuates bacterial growth by inhibiting L-serine metabolism(Cosloy and McFall, 1973). Therefore, bacterial sensitivity to D-serine may reveal a common pathway for inhibition of mitochondrial one-carbon metabolism by D-serine. Given that neurons do not express enzymes for mitochondrial one-carbon metabolism upon maturation (Figure 4 –figure supplement 9), evolutionary acquisition of neurophysiological functions of D-serine without cytotoxicity may be due to the fact that mature neurons are not proliferative and do not depend on this conserved metabolic feature of mitochondria. While complete understanding of the significance of serine stereo-conversion in the developmental brain will require further studies to illuminate neurogenesis under spatio-temporally controlled D-serine, our findings provide a basis to understand the significance of maintaining serine enantiomer balance in the central nervous system.

## Materials and Methods

| **Key Resources Table** |  |  |  |  |
| --- | --- | --- | --- | --- |
| **Reagent type (species) or resource** | **Designation** | **Source or reference** | **Identifiers** | **Additional information** |
| gene (include species here) | Human SHMT2 | This study |  |  |
| strain, strain background (Mus musculus) | C57BL/6Jcl | CLEA Japan (Tokyo, Japan) | MGI:3055581 |  |
| strain, strain background (Mus musculus) | CrI:CD1(ICR) | Charles River Laboratories (Wilmington, MA, USA) | MGI:5649524 |  |
| strain, strain background (Mus musculus) | BALB/c nude mice (BALB/cSlc-nu/nu) | Japan SLC |  |  |
| cell line (mouse) | Neuro2a | ATCC | CCL-131 |  |
| cell line (mouse + rat) | F11 | Platika D et al. (PMID: 3858835) |  |  |
| cell line (human) | SHSY5Y | ATCC | CRL-2266 |  |
| cell line (mouse) | NSC34 | Cashman NR et al., (PMID: 1467557) |  |  |
| cell line (human, male) | U87 | ATCC | HTB-14 |  |
| cell line (human, male) | U251 | JCRB Cell Bank | IFO50288 |  |
| cell line<br>(human) | SF126 | JCRB Cell Bank | IFO50286 |  |
| cell line<br>(human) | hG008 | Fukaya R et al.<br>(PMID:<br>26980763) |  |  |
| cell line<br>(human) | hG020 | Fukaya R et al.<br>(PMID:<br>26980763) |  |  |
| transfected<br>construct | Sfxn1-targeting<br>siRNA | Dharmacon | ON-TARGETplus<br>Mouse Sfxn1<br>(14057) siRNA<br>SMARTpool |  |
| transfected<br>construct | non-targeting<br>control<br>siRNA | Dharmacon | ON-TARGETplus<br>Non-targeting<br>siRNA#1 |  |
| biological<br>sample (*Mus*<br>*musculus*) | Primary<br>cortical<br>neurons | This paper |  | Freshly<br>isolated<br>from<br>embryonic<br>day 14 *Mus*<br>*musculus* |
| biological<br>sample (*Mus*<br>*musculus*) | Primary<br>cortical<br>astrocytes | This paper |  | Freshly<br>isolated<br>from<br>embryonic<br>day 14 *Mus*<br>*musculus* |
| antibody | Cleaved<br>Caspase-3<br>(Asp175)<br>(5A1E)<br>Rabbit<br>Monoclonal<br>Antibody | Cell Signaling<br>Technology | Cat# 9664<br>RRID:AB_2070042 | WB 1:1000<br>IF 1:200 |
| antibody | Rabbit anti-GAPDH,<br>clone 14C10 | Cell Signaling<br>Technology | Cat# 2118<br>RRID:AB_561053 | WB 1:2000 |
| antibody | Rabbit anti-GFAP | Agilent | Cat# Z0334<br>RRID:AB_10013382 | IF 1:200 |
| antibody | Mouse anti-vGlut2, clone 8G9.2 | Abcam | Cat# Ab79157<br>RRID:AB_1603114 | IF 1:200 |
| antibody | Mouse anti-GAD67, clone 1G10.2 | Merck-Millipore | Cat# MAB5406<br>RRID:AB_2278725 | IF 1:200 |
| sequence-based reagent | Rabbit Anti-Sfxn1 polyclonal antibody | Sigma-Aldrich | Cat# HPA019543<br>RRID:AB_1856789 |  |
| Recombinant DNA reagent | pET41a vector | Novagen |  |  |
| peptide, recombinant protein | human SHM T2 | This study |  |  |
| commercial assay or kit | EnzyChrom Formate Assay Kit | BioAssay Systems | Catalog No: EFOR-100 |  |
| software, algorithm | MetaboAnalyst | https://www.metaboanalyst.ca/ | RRID:SCR_015539 |  |
| software, algorithm | Trim Galore | https://www.bioinformatics.babraham.ac.uk/projects/trim_galore/ | RRID:SCR_011847 |  |
| software, algorithm | HISAT2 | https://daehwankimlab.github.io/hisat2/ | RRID:SCR_015530 |  |
| software,<br>algorithm | DESeq2 | https://bioconductor.org/packages/release/bioc/html/DESeq2.html | RRID:SCR_015687 |  |
| software,<br>algorithm | ClusterProfiler | https://bioconductor.org/packages/release/bioc/html/clusterProfiler.html | RRID:SCR_016884 |  |

### Animals

All animal experiments were approved by the institutional Animal Experiment Committee and conducted in accordance with Institutional Guidelines on Animal Experimentation at Keio University. C57BL/6Jjcl mice were purchased from CLEA Japan (Tokyo, Japan). ICR mice were from Charles River Laboratories (Wilmington, MA, USA). Female BALB/c nude mice (BALB/cSlc-nu/nu) were from Japan SLC (Shizuoka, Japan). Mice were raised in 12 h light and dark cycle with free access to food (CE-2, CLEA Japan) and water in the specific pathogen-free environment.

### Antibodies

Rabbit polyclonal antibodies to cleaved caspase-3 and to GAPDH were purchased from Cell Signaling Technologies (Danvers, MA, USA). Mouse monoclonal antibodies to GFAP, vGLUT2, and GAD67 were from Agilent Technologies (Santa Clara, CA, USA), Abcam (Cambridge, UK), and Merck Millipore (Darmstadt, Germany), respectively.

### Cell culture and proliferation assay

NPCs and primary cultured astrocytes (PCAs) were prepared as follows. Telencephalon tissues from ICR mice were dissected at the embryonic day 14, treated with papain, and disassembled by pipetting. The cell suspension was passed through a 40-μm filter, centrifuged, and resuspended in a Neurobasal medium without serine (NB-S). NB-S was manufactured by Research Institute for the Functional Peptide (Yamagata, Japan) based on the concentration of each chemicals in Neurobasal media (ThermoFisher: 21103049) described in the technical resources at ThermoFisher. NPCs were cultured in NB-S supplemented with 2% B-27 supplement (ThermoFisher, Waltham, MA, USA), 1% Penicillin-streptomycin solution (ThermoFisher), and 1% GlutaMAX supplement (ThermoFisher). For PCAs, NPCs were differentiated into astrocytes within 7 days by changing the medium to D-MEM (high glucose) (Fujifilm Wako, Tokyo, Japan) supplemented with 10% FBS (A5256701, ThermoFisher) and 1% Penicillin-Streptomycin at 24 h *in vitro*.

Neuroblastoma (Neuro2a, F11, NSC34, and SHSY5Y) and glioma (U87, U251, SF126) cell lines were cultured in MEM supplemented with 10% dialyzed FBS (S-FBS-NL-065, SERENA Europe GmbH, Brandenburg, Germany), 5 mM D-glucose, 65 μM sodium pyruvate, 1 × MEM vitamin solution (ThermoFisher), 2 mM L-glutamine, 0.15 mM L-proline, 0.15 mM L-alanine, 0.15 mM L-aspartic acid, 0.15 mM L-glutamic acid, and 0.34 mM L-asparagine (-SG media) as described in Tajan et al.(Tajan et al., 2021), unless otherwise noted. Proliferation of NPCs, PCAs, and cancer cell lines was analyzed with Cell Counting Kit-8 in accordance with the manufacture’s protocol (Dojindo, Kumamoto, Japan).

### Western blot

NPCs were harvested in a lysis buffer [50 mM Tris-HCl, pH 7.4, 150 mM NaCl, 20 mM ethylenediaminetetraacetate (EDTA), 1% TritonX-100, protease inhibitor cocktail (Sigma-Aldrich, St. Louis, MO, USA)]. Protein concentration was analyzed with a BCA assay kit (ThermoFisher), and 20-80 µg of protein were subjected to SDS-PAGE. Then, proteins were transferred to the PVDF membranes and blocked with 10% skim milk in PBST. Proteins were detected with anti-cleaved caspase-3 antibody (clone: 5A1E) (Cell Signaling Technology, MA, USA), anti-Sfxn1 antibody (HPA019543) (Sigma-Aldrich), or ant-GAPDH antibody (clone: 14C10) (Cell Signaling Technology), and HRP-conjugated secondary antibodies (Jackson ImmunoResearch Laboratories, PA, USA).

### Metabolomics

Cells were washed twice with 10 mL of PBS(-). The PBS(-) was aspirated, and 1 mL of MeOH containing internal standards was added to the dish. The dish was left for 10 min, and a sample solution was obtained and transferred to a tube. For metabolite extraction, the tube was vortexed and subsequently centrifuged at 20,380×g for 10 minutes at 4 °C (MDX-310, TOMY Seiko, Tokyo, Japan). The 150 μL of supernatant was transferred to another tube and dried by centrifugation at 1,600 rpm (366×g) for 90 min at room temperature (VC-96W, Taitec, Saitama, Japan). Then, 10 μL of 90% MeOH were added, 30 μL of H_2_O were added and mixed, and then the tube was centrifuged at 20,380×g for 10 minutes at 4 °C. The 20 μL of supernatant were transferred to a vial and injected into the LC-MS system.

The LC-MS instrument has previously been described in detail(Fuse et al., 2020). Briefly, an Agilent Technologies 1290 Infinity LC system and a G6230B time-of-flight MS (TOF–MS) (Agilent Technologies, Santa Clara, CA, USA) were used. Each sample was analyzed in positive and negative modes. Conditions for the analysis in positive mode were set as described previously, with slight modification(Tomita et al., 2018). The temperature of the LC columns was set at 40 °C. For negative mode, the chromatographic separation was performed using an ACQUITY HSS T3 column (2.1 i.d. × 50 mm, 1.8 μm; Waters, Milford, MA, USA) at 30 °C. The mobile phase, consisting of solvent A (0.1% formic acid in water) and solvent B (acetonitrile), was delivered at a flow rate of 0.3 mL/min. In this study, 50–1,200 m/z was used for the MS setting in negative mode.

Data processing from raw LC-MS data to produce a data matrix including metabolite concentration (sample × metabolite) was described previously(Pang et al., 2021). Data analyses were performed using MetaboAnalyst (ver. 5.0, https://www.metaboanalyst.ca/)(Pang et al., 2021) to produce volcano plots and to conduct pathway analysis and pattern searches. Overall data profiles were visualized using MeV TM4 (ver. 4.9.0, https://sourceforge.net/projects/mev-tm4/).

### Quantification of glycine and serine enantiomers

Glycine and serine enantiomers in cell-cultured media or brain tissues were measured using two-dimensional HPLC, as previously described(Gonda et al., 2023; Ishii et al., 2018). Briefly, samples were mixed with 9 volumes of MeOH and centrifuged to remove protein depositions. Supernatants were spin-dried and resuspended in Milli-Q water. Resuspensions were resuspended in 200 mM sodium borate, and then derivatized with 4-fluoro-7-nitro-2,1,3-benzoxadiazole (NBD-F). NBD-conjugated amino acids were injected into a two-dimensional HPLC system (NANOSPACE SI-2 series, Shiseido, Tokyo, Japan), separated on an octadecylsilyl column (Singularity RP18, 1.0 mm inner diameter (ID) × 250 mm) (designed by Kyushu University and KAGAMI Co. Ltd., Osaka, Japan), and further separated into enantiomers on a Pirkle-type enantioselective column (Singularity CSP-001S, 1.5 mm ID × 250 mm) (designed by Kyushu University and KAGAMI). Fluorescence of NBD-amino acids was detected at 530 nm with excitation at 470 nm. D- and L-serine peak in a chromatogram were detected and standard curve was drawn by the peak height of those standards. The absolute amount of D- and L-serine were calculated by standard curve method.

### Quantification of formate in the culture media

Formate concentration in the culture media was measured using Enzychrom Formate assay kit (BioAssay Systems) according to the manufacturer’s protocol.

### SHMT2 activity assays

The SHMT2 gene was cloned from cDNA of HeLa cells using primers 5’-GCCCATATGGCCATTCGGGCTCAGCAC-3’ (forward) and 5’-GCCCTCGAGATGCTCATCAAAACCAGGCA-3’ (reverse) and subcloned into the pET41a vector (Novagen, WI, USA) at restriction enzyme sites of NdeI and XhoI. C-terminally hexahistidine (His_6_)-tagged human SHMT2 was purified as follows. *E. coli* BL21 (DE3) pLysS-competent cells were transformed with pET41-SHMT2, cultured at 37 °C in Luria-Bertani medium containing 20 µg/mL kanamycin, and grown until the OD600 reached 0.6-0.7. Isopropyl β-D-thiogalactopyranoside was added to a final concentration of 0.5 mM, and culturing was continued for an additional 24 h at 20°C. Harvested cells were resuspended in 20 mM sodium phosphate buffer (pH7.4) containing 500 mM NaCl and protease inhibitors (Nacalai Tesque, Kyoto, Japan), and sonicated using a Sonifier 250 instrument (Branson, CT, USA). Lysed cells were centrifuged at 20,000 × g for 10 min at 4 °C, and imidazole was added to the supernatant to a final concentration of 100 mM. The supernatant was applied to a His SpinTrap column (Cytiva, Tokyo, Japan). The column was washed with 20 mM sodium phosphate buffer (pH7.4) containing 500 mM NaCl and 100 mM imidazole. and recombinant protein was eluted with 20 mM sodium phosphate buffer (pH7.4) containing 500 mM NaCl and 500 mM imidazole. Protein fractions were buffer-exchanged into 50 mM Tris-HCl (pH8.0) containing 100 mM NaCl and 10% glycerol using an Amicon Ultra-0.5 centrifugal filter 10K device (Merck Millipore, Darmstadt, Germany).

SHMT2 activity was determined by measuring the production of glycine. The reaction mixture (150 µL) contained 50 mM HEPES-NaOH (pH8.0), 0.2 mM L-serine, 50 μM PLP, 50 μM THF, 1 mM dithiothreitol, and recombinant SHMT2 (2 µg). D-serine was added at concentrations varying from 0.1 to 0.4 mM. The reaction mixture was incubated at 37 °C for 10 min, and then 600 µL of MeOH were added to terminate the reaction. Glycine was derivatized with ortho-phthaldialdehyde (OPA) and *N*-tert-butyloxycarbonyl-L-cysteine (Boc-L-Cys), and an aliquot (10 µL) was injected into an LC-4000 Series HPLC system (Jasco Corp., Tokyo, Japan) equipped with a Mightysil RP-18GP column (150 × 4.6 mm i.d.; Kanto Chemical Co., Tokyo, Japan). A gradient of solvent A (50 mM sodium acetate buffer, pH 6.0) and solvent B (acetonitrile) was applied at a flow rate of 1 mL/min. A linear gradient was applied for 35 min from 10% to 21% solvent B for glycine analysis. Excitation and emission wavelengths for fluorescence detection were 344 nm and 443 nm, respectively.

### L-serine transport assay in semi-permeabilized cells

Neuro2a cells were were seeded on poly-D-lysine coated 24-well-plates and transfected with either Sfxn1-targeting siRNA (ON-TARGETplus Mouse Sfxn1 (14057) siRNA SMARTpool, Dharmacon) or non-targeting control siRNA (ON-TARGETplus Non-targeting siRNA#1, Dharmacon) using Lipofectamine 2000 (Thermo Fisher Scientific) according to the manufacturer’s instructions. Cells were cultured for 3 days after transfection and then subjected to the transport assay. Transport of L-[^3^H]serine was examined in semi-permeabilized cells. For plasma membrane permeabilization, cells were treated with saponin, an agent that permeabilizes the plasma membrane but leaves membranous organelles intact(Klepinin et al., 2016; Kuznetsov et al., 2008). The optimized saponin concentration and treatment time were verified by optical microscopy. Transport assays were performed as described previously(Lee et al., 2022; Wiriyasermkul et al., 2012) with some modifications. After washing the cells with 37°C pre-warmed Dulbecco’s Phosphate Buffered Saline (DPBS) with 1 mM MgCl_2_ and 0.25 mM CaCl_2_, they were incubated for 6 min in the same buffer containing 40 μg/mL saponin (Fujifilm) for plasma membrane permeabilization. Transport of 10 μM L-[^3^H]serine (100 Ci/mol; Moravek Biochemicals, Brea, CA, USA) with or without D-serine (at indicated concentrations) was measured for 10 min in the same buffer including saponin. After terminating the reaction and cell lysis, an aliquot of the lysate was used to measure protein concentration by BCA protein assay (Takara Bio). The lysate was mixed with OptiPhase HiSafe 3 (PerkinElmer), and radioisotope activity was monitored by LSC-8000 β-scintillation (Hitachi, Tokyo, Japan). Data shown in the figures were those subtracted from uptake values at 0 min. Kinetics of inhibition was fitted to the Nonlinear Regression: dose-response – inhibition model of GraphPad Prism 8.4.

### RNA-sequence and analysis

Neuro2a cells were cultured in D-MEM with 10% FBS. The medium was changed to D-MEM(-SG) including 10% dialyzed FBS with or without 5 mM D-serine. Cells were harvested in Trizol after 18 h of incubation at 37 °C in a CO_2_ incubator. RNA was extracted from cells according to the manufacture’s protocol. mRNA libraries of each sample were prepared using a TruSeq Stranded mRNA library Kit (Illumina, San Diego, CA) according to the manufacturer’s protocol and paired end sequences were read by NovaSeq (Illumina). Adapter sequences were trimmed from raw sequences with Trim Galore (https://www.bioinformatics.babraham.ac.uk/projects/trim_galore/) and trimmed sequences were mapped to the mouse genome (GRCm38/mm10) using HISAT2 (https://daehwankimlab.github.io/hisat2/). Aligned sequences were counted using FeatureCounts (https://subread.sourceforge.net/). Differentially expressed genes were identified by DESeq2 (https://bioconductor.org/packages/release/bioc/html/DESeq2.html). Gene Set Enrichment analysis was performed to find enriched biological pathways using ClusterProfiler (https://bioconductor.org/packages/release/bioc/html/clusterProfiler.html) with the following settings: pAdjustMethod = fd, pvalueCutoff = 0.05, minGSSize = 10, maxGSSize = 400.

### Glioma stem cell culture

hG008 and hG020 cells, established from human glioblastoma specimens(Fukaya et al., 2016), were cultured in Ultra-Low attachment cell culture flasks (Corning, Kennebunk, ME, USA) in a medium, in which ingredients were manually mixed in MEM to make DMEM/Ham’s F-12 with HEPES without serine and glycine (D/H-SG). The D/H-SG medium was supplemented with 2% B-27 (ThermoFisher), 20 ng/mL recombinant human fibroblast growth factor-basic (PeproTech, Rocky Hill, NJ, USA), 20 ng/mL recombinant human epidermal growth factor (PeproTech), 1000 units/mL recombinant human leukemia inhibitory factor (Nacalai Tesque, Kyoto, Japan), and 1 unit/mL heparin. Sphere formation of hG008 cells (1×10^4^ cells/well in 96-well plates with ultra-low attachment surface, Corning) was observed using EVOS M5000 (ThemoFisher) and the maximum diameter of spheres in each well was measured after 7 days of culture.

### Ex vivo cancer growth assay

U87 and hG008 cells were transduced with the lentiviral vector CSII-EF-ffLuc, and single-cell suspensions were cultured in 96-well plates. Xenografts were implanted in the brain of female BALB/c nude mice as described previously^25^. Briefly, mice were implanted with Venus fluorescence-labeled U87 (5×10^5^ cells/2 μL in PBS) or hG008 (1×10^5^ cells/2 μL in PBS) under anesthesia on a stereotaxic apparatus (Narishige Scientific Instrument Lab, Tokyo, Japan). Mice were sacrificed after 6 days (U87) or 35 days (hG008) of implantation. Mouse brains were dissected, cut into a 200-μm slices, and cultured on the Millicell cell culture inserts (Millipore-Sigma, Burlington, MA) placed on a 35 mm glass-bottom dish filled with D/H-SG medium with or without 100 mM D-serine. Fluorescence images were taken with a FV3000 (Olympus, Tokyo, Japan) on days 0 and 7 and analyzed with Image J software (https://imagej.nih.gov/ij/index.html).

### Statistics

No statistical methods were used to predetermine sample size. Blinding was not performed. No randomization was used. Prism 10 (GraphPad Software) was used for data plotting and statistical analyses. Statistical significance was determined with unpaired t-tests to compare two groups, or one-way analysis of variance (ANOVA) for multiple comparisons when data were normally distributed and had equal variance. ANOVA was followed by Dunnett’s post-hoc test (comparison to control) or Tukey’s post-hoc test (comparisons among samples).

## Data Availability

RNA-seq data are available at Gene Expression Omnibus (GEO) with accession number GSE268106.

## Acknowledgements

We thank Kenji Hamase and Masashi Mita for technical support on chiral amino acid analysis and Steven D Aird for editing the manuscript. This study was supported by the Grants-in-Aid for Scientific Research (KAKENHI) 15K19497 (MS), 17J10213 (MS), 22K19408 (JS), Keio Gijuku Fukuzawa Memorial Fund for the Advancement of Education and Research (JS), and Keio Program for the Promotion of Next Generation Research Projects Type A (JS).

## Supplemental Information

### Materials and Methods

#### Preparation of model systems for MD simulations

Model systems of SHMT2 combined with either L- or D-serine in the presence of THF and PLP were prepared for molecular dynamics (MD) simulations. The X-ray crystallographic structure of SHMT2 (PDB ID: 4PVF)(Giardina et al., 2015)was obtained from the protein data bank (PDB)(Berman et al., 2000) and removed small molecules including crystal water molecules. Missing atoms and hydrogen atoms in the SHMT2 structure were added by using homology modeling techniques with the Molecular Operating Environment (MOE) program (Chemical Computing Group ULC, 910–1010 Sherbrooke St. W. Montreal, Quebec, H3A 2R7, Canada. https://www.chemcomp.com/en/index.htm). The L- or D-serine-bound SHMT2 complex structures were constructed by placing serine and THF molecules referring from the X-ray crystallographic structure of *At*SHMT2-MTX complex ((PDB ID: 6SMN))(Ruszkowski et al., 2019). Then, these complex structures were solvated in a rectangular box (130 × 137 × 155Å^3^) containing 0.15 M NaCl in TIP3P water molecules. Each model system contains 30,136 atoms.

#### Molecular dynamics simulations

All MD simulations were performed using the GROMACS version 5.0.6 program(Abraham et al., 2015; Berendsen et al., 1995). The AMBER14SB force field(Maier et al., 2015) was used for protein and general amber force field (GAFF)(Wang et al., 2004) was used for THA and PLP. The partial charges for the ligands were calculated at the RHF/6-31G(d) level with Gaussian 03(Frisch et al., 2004) and the restrained electrostatic potential method. The periodic boundary condition was applied to the initial system, and the temperature and pressure were kept constant using the Nosè-Hoover thermostat(Shūichi Nosé, 1984; Shuichi Nosé, 1984) and Parrinello-Rahman barostat(Parrinello and Rahman, 1981), respectively. The linear constraint solver (LINCS) algorithm(Hess et al., 1997) was applied to all covalent bonds, with an integration time step of 2.0 fs taken into consideration. The long-range Coulomb interactions were treated with the particle mesh Ewald method(Essmann et al., 1995) and the direct space cut-off distance was set to 10 Å. The van der Waals interactions were calculated using a switched cut-off between 8 and 10 Å. After the energy minimization of each solvated system, the system was gradually heated to 300 K during 100 ps. Then, 2 × 300 ns MD simulations under NPT ensemble (P = 1 bar and T = 300 K) were performed for L- or D-serine-SHMT2 complex model systems.

#### Immunocytochemistry

NPCs were cultured on cover glasses coated with poly-L-lysine. After treatment with D-serine, NPCs were fixed in 4% paraformaldehyde PBS at 4 ℃ for more than 4 h, rinsed in PBS, and permeabilized/blocked in PBS containing 5% goat serum and 0.3% TritonX-100 for 1 h. Then, cells were co-immunolabeled with primary antibodies to Nestin, beta III tubulin isoform b, vGlut2, or GAD67 and a rabbit polyclonal antibody to cleaved caspase-3 (Cell Signaling Technology, Danvers, MA, USA) in 5% goat serum PBS, and were further immersed in PBS containing a goat anti-rabbit IgG FITC-conjugate antibody and a goat anti-mouse IgG Texas Red-conjugated antibody with 5% goat serum. After rinsing 2 times in PBS, cells were coverslipped in ProLong Gold Antifade reagent with DAPI (Life Technologies, Carlsbad, CA). Cells were observed with a confocal microscope (LSM310, Zeiss, Germany).

#### Quantification of glutathione in NPCs

NPCs (8×10^5^ cells/6-well plate) were treated with D- or L-serine for 48 hours. Cells were washed twice with 2 mL of PBS and harvested in 80 mL of 10 mM HCl using a cell scraper. Glutathione concentration in the cell extract was measured using the GSSG/GSH quantification kit II (Dojindo, Kumamoto, Japan) according to the manufacturer’s protocol. GSSG was not detected in our samples. Protein concentrations in those samples were measured by the BCA assay and used to normalize GSH concentration.

#### Lipidomics

NPCs (4×10^6^ cells/ 10 cm dish) were cultured in NB-S (serine-free Neurobasal medium, Research Institute for the Functional Peptide) and treated with 2 mM D-serine at DIV2 for 48h. For controls, NPCs were treated with vehicle. Three biological replicates were washed twice in PBS, harvested in 100% ethanol, and mixed into one sample for each group.

The lipid composition of NPCs was analyzed with liquid chromatography-mass spectrometry (LC-MS). Cell phospholipids extracted in 100% ethanol were evaporated, resuspended in methanol, and mixed with chloroform. After separation by centrifuge, the supernatant mixed with internal control (Phosphatidylcholine, 16:0 D30/18:1) was subject to LC-MS analysis. Lipids in samples were separated on an L-column2 ODS metal free column (50 mm, × 2 mm I.D., particle size 3 μm, CERI) in the mobile phase (A) 1 mM formate, (B) 1mM formate in 2-propanol, and (C) acetonitrile at 0.3 mL/min, and analyzed by electrospray ionization using an LTQ Orbitrap XL spectrometer (Thermo Fisher Scientific) with Data-dependent acquisition of MS/MS spectra. LipidSearch4.1 software (Mitsui Knowledge Industry, Japan) and the Orbitrap database were used for peak picking, lipid molecule estimation, and sample alignment.

#### Analysis of phosphatidyl-D/L-serine

Lipids were extracted from NPCs using the Bligh and Dyer methods. Briefly, cells were scraped and suspended in 50 μL of water. Cell suspensions were mixed with 187.5 μL of chloroform: methanol (1:2), vortexed briefly, and added to 62.5 μL of chloroform and 62.5 μL of water. After centrifugation at 3,300 × *g* for 10 min, the lower layer was collected, washed in water: methanol: chloroform (47:48:3), and dried. Pellets of lipids were resuspended in a phospholipase D (PLD) reaction buffer [50 mM NaCl, 50 mM Tris-HCl (pH 8.0), 1 mM CaCl_2_, 0.1% Triton X-100, 1 U/μL PLD] and incubated at 37 ℃ for 1 h. After the reaction with PLD, samples were mixed with methanol to bring them up to 80% methanol and centrifuged at 20,400 × *g* for 10 min. The supernatant was dried up, and the remaining residue was resuspended in 200 mM borate. D- and L-serine from phosphatidyl-serine were analyzed using the 2D HPLC.

#### Public RNA-seq data analysis

Bulk RNA-seq count data of ES cell-derived neuron(Hubbard et al., 2013) were downloaded from http://dx.doi.org/10.6084/m9.figshare.154970, and those of iPSC-derived cortical neuron(Burke et al., 2020) were from http://stemcell.libd.org/scb/data_links.html. Relative expression levels among developmental stages were illustrated as z-scores of reads per kilobase million (RPKM).

Single-cell RNA-seq data of mouse cerebral cortex were downloaded from Gene Expression Omnibus (GSE153164) and analyzed using Seurat as described previously(Bella et al., 2021) with small modifications. Briefly, all datasets from E10 to P4 were merged (technical duplicate of P1 was not integrated, in order to simplify the analytical pipeline) and filtered by cut-off values of nFeature_RNA > 200, nCount_RNA > 500, and percent.mt < 7.5. The filtered dataset was normalized using the SCTransform function with the “variance stabilizing transformation” method, and integrated using SelectIntegrationFeatures (nfeatures = 3000), PrepSCTIntegration, and FindIntegrationAnchors with the SCT normalization method. After integration, data dimensions were reduced using RunPCA. Clustering was performed using FindNeughbors with PCA reduction and FindClusters with the resolution setting 0.6. Uniform manifold approximation and projection (UMAP) were calculated with RunUMAP (reduction = pca, metric = euclidean). Clusters of cells were generated with Dimplot and gene expression profiles were visualized with Featureplot.

### Supplemental Figures

**Figure 1 figure supplement 1.**
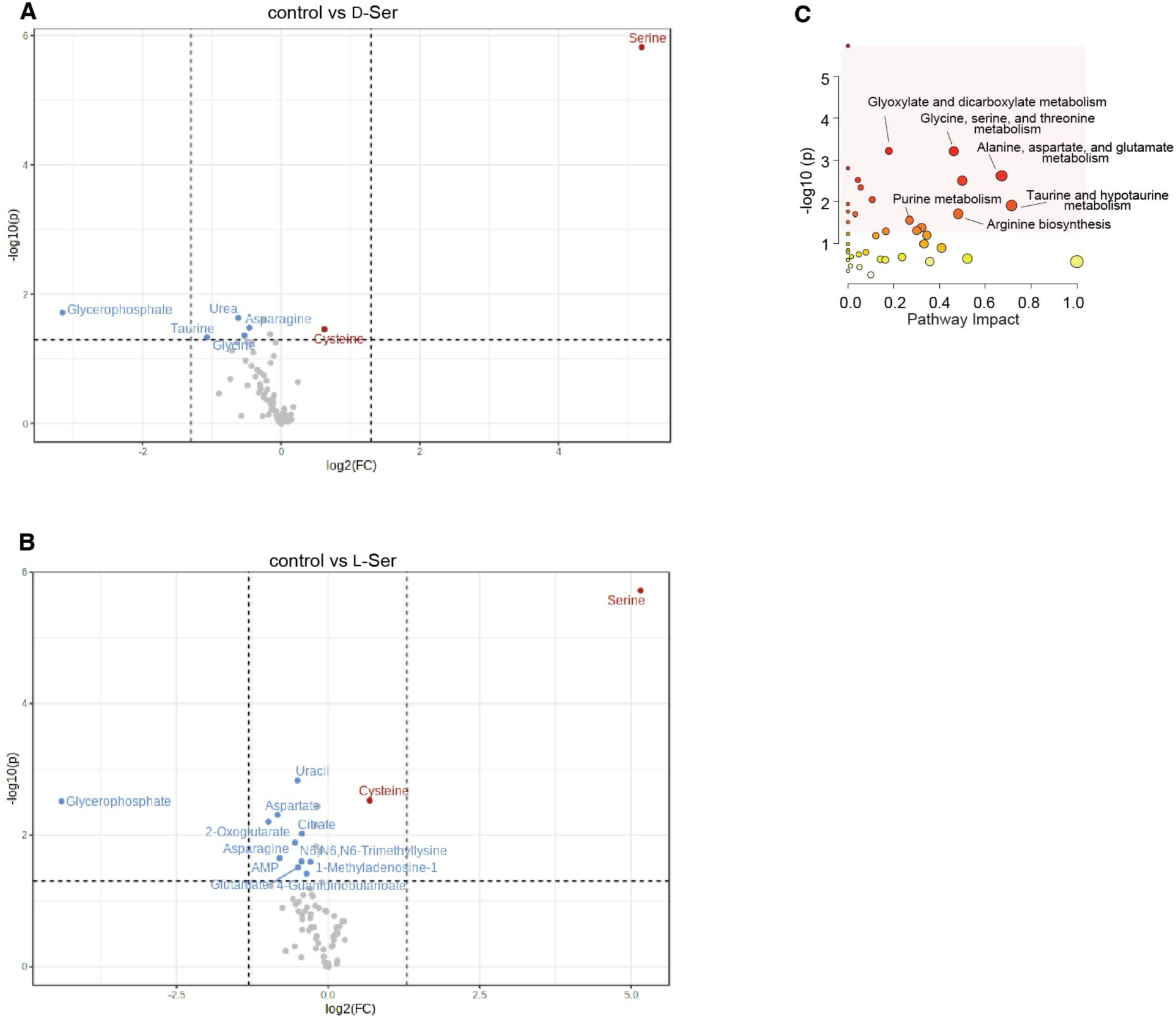
Metabolomic profiles of PCNs treated with D- or L-serine A, **B**. PCNs were treated with vehicle-control, L-serine, or D-serine, and processed for metabolomic analysis. Volcano plots show differentially detected metabolites by comparing control vs D-serine (**A**) or L-serine (**B**). Horizontal axis, log_2_ foldchange. Vertical axis, –log_10_(p-value). Increased metabolites, red. Decreased, blue. **C**. Impact of metabolite differences detected between vehicle and L-serine in KEGG metabolic pathways. Horizontal axis, the pathway impact. Vertical axis, –log_10_(p-value) of each pathway.

**Figure 2 figure supplement 2.**
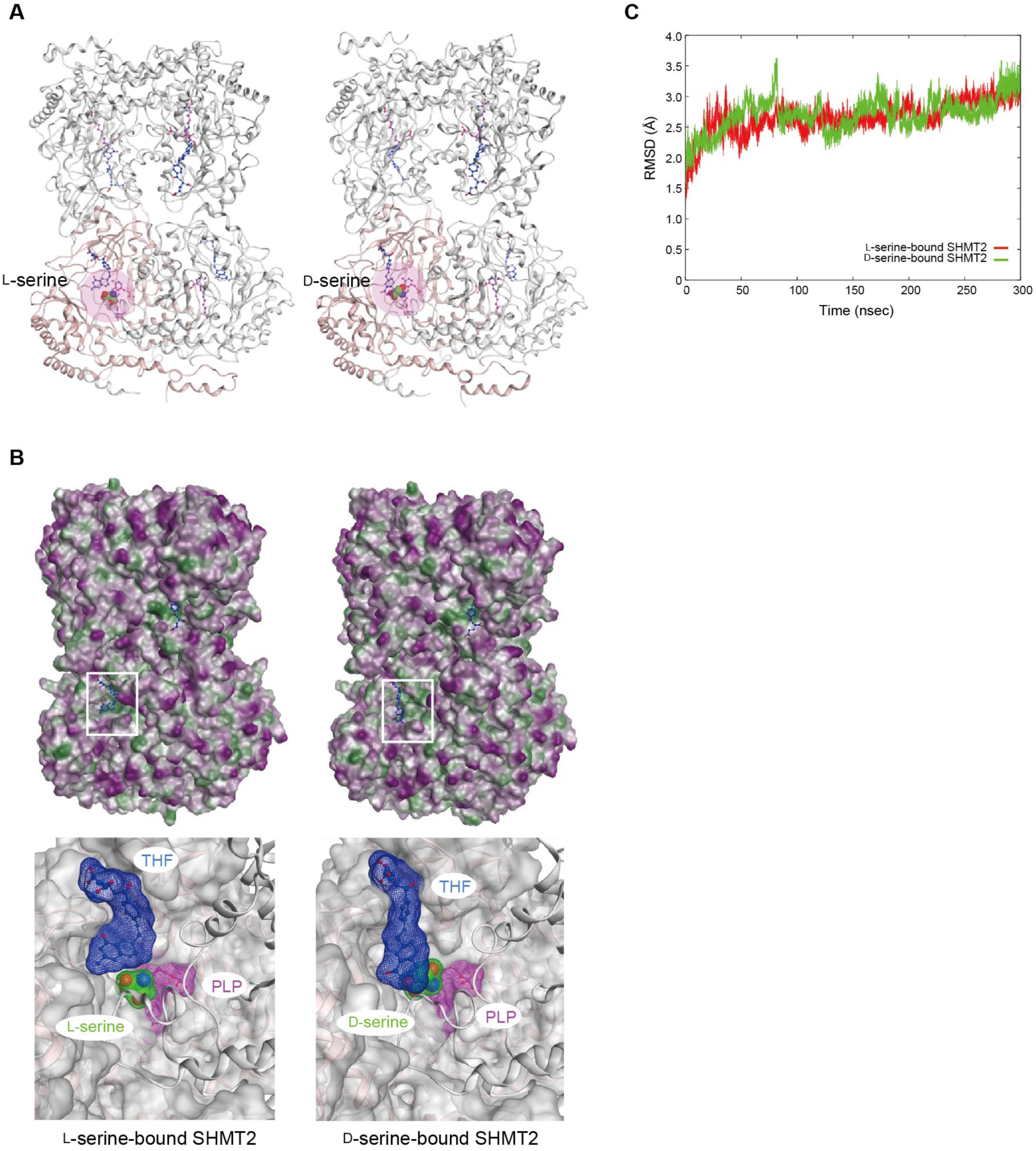
Molecular dynamics simulations for serine-enantiomer-bound SHMT2. **A**. Ribbon form of the initial structures of L- or D-serine-bound SHMT2. L- and D-serine show space filling models. **B**. (Top) Molecular Surface of the initial structures of L- or D-serine-bound SHMT2. Purple, hydrophilic parts. Green, lipophilic parts. (Bottom) Close-up views of the binding pockets of L-serine (left) and D-serine (right). Purple, PLP. Blue, THF. Green, serine. **C**. The root mean square deviation (RMSD) values of L-serine (red) or D-serine (green)-bound SHMT2 during molecular dynamics (MD) simulations.

**Figure 2 figure supplement 3.**
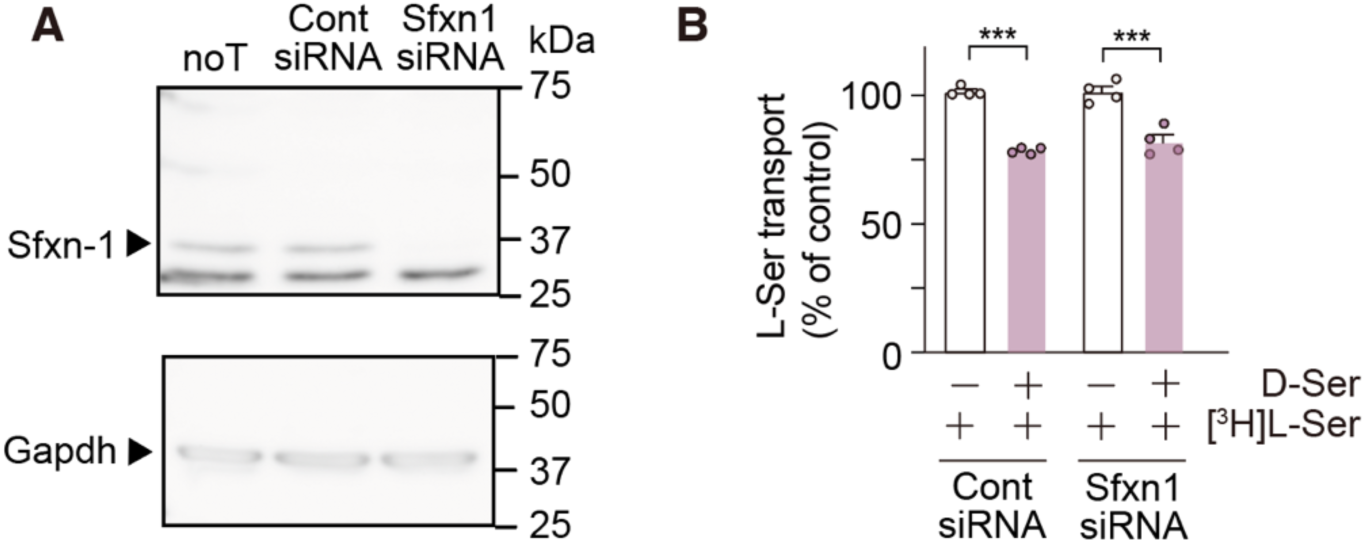
Effect of Sfxn1 knockdown on D-serine-mediated inhibition of mitochondrial L-serine transport. **A**. Western blots indicate Sfxn1 and Gapdh in Neuro2a cells transfected with or without Sfxn1-targeting or control siRNA. noT, non-transfected. cont, control. **B**. Inhibition of mitochondrial L-serine transport by 1 mM D-serine was examined in semi-permeabilized Neuro2a cells transfected with Sfxn1-targeting or control siRNA (n = 4, each). Statistical significance was analyzed by one-way ANOVA. *** *p* < 0.001.

**Figure 2 figure supplement 4.**
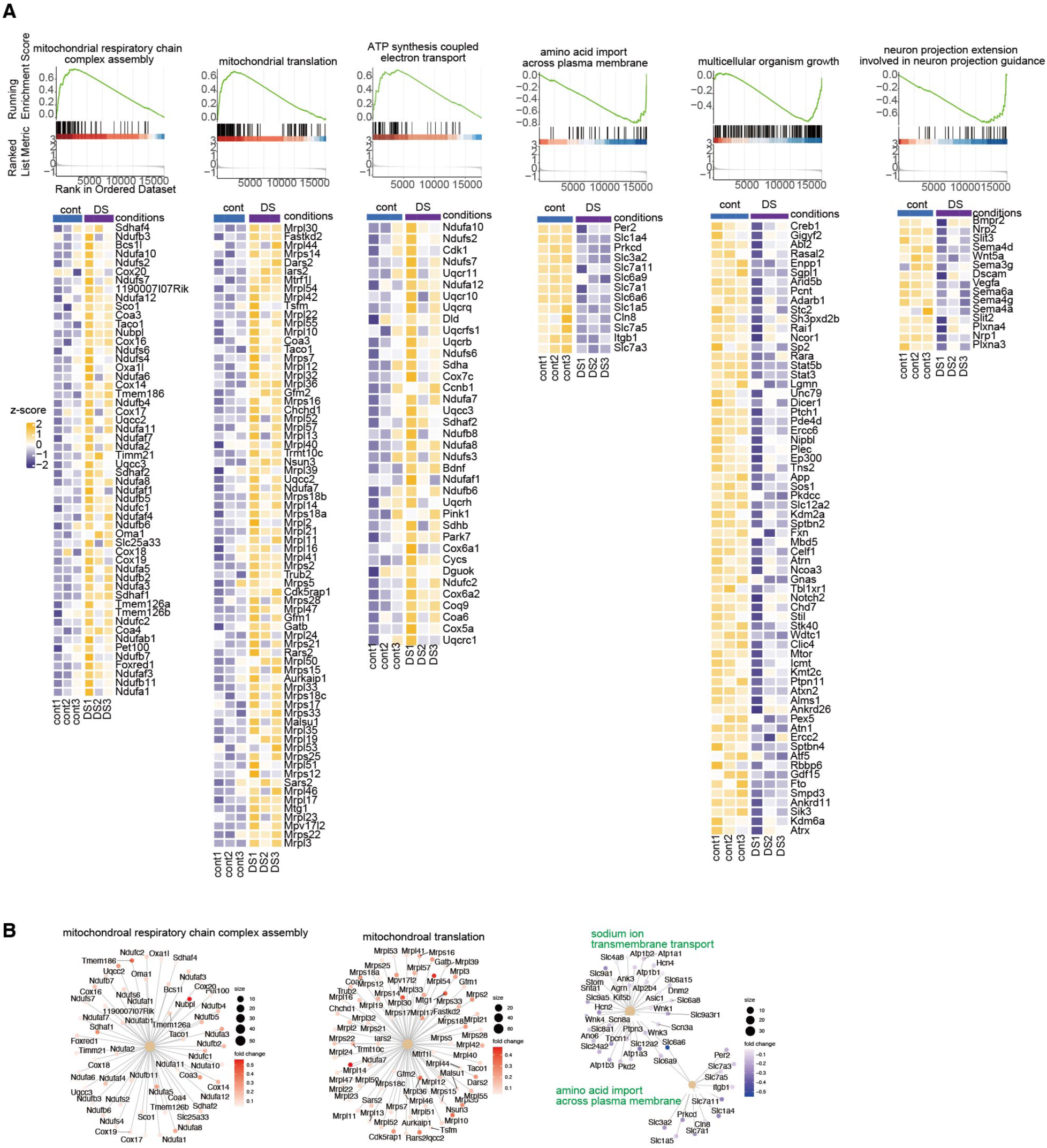
Gene sets up- or down-regulated by D-serine in Neuro2a cells. **A.** RNA-seq was performed using total RNA extracted from Neuro2a cells treated with vehicle-control or D-serine (DS). Upper panels show GSEA plots of ontologies altered by D-serine treatment. Bottom heatmaps present z-scores of gene expressions significantly altered by D-serine, by gene ontology. **B.** Network plots indicate up- or down-regulated genes categorized by ontology. Color represents fold-change expressions in D-serine treated samples compared to controls.

**Figure 3 figure supplement 5.**
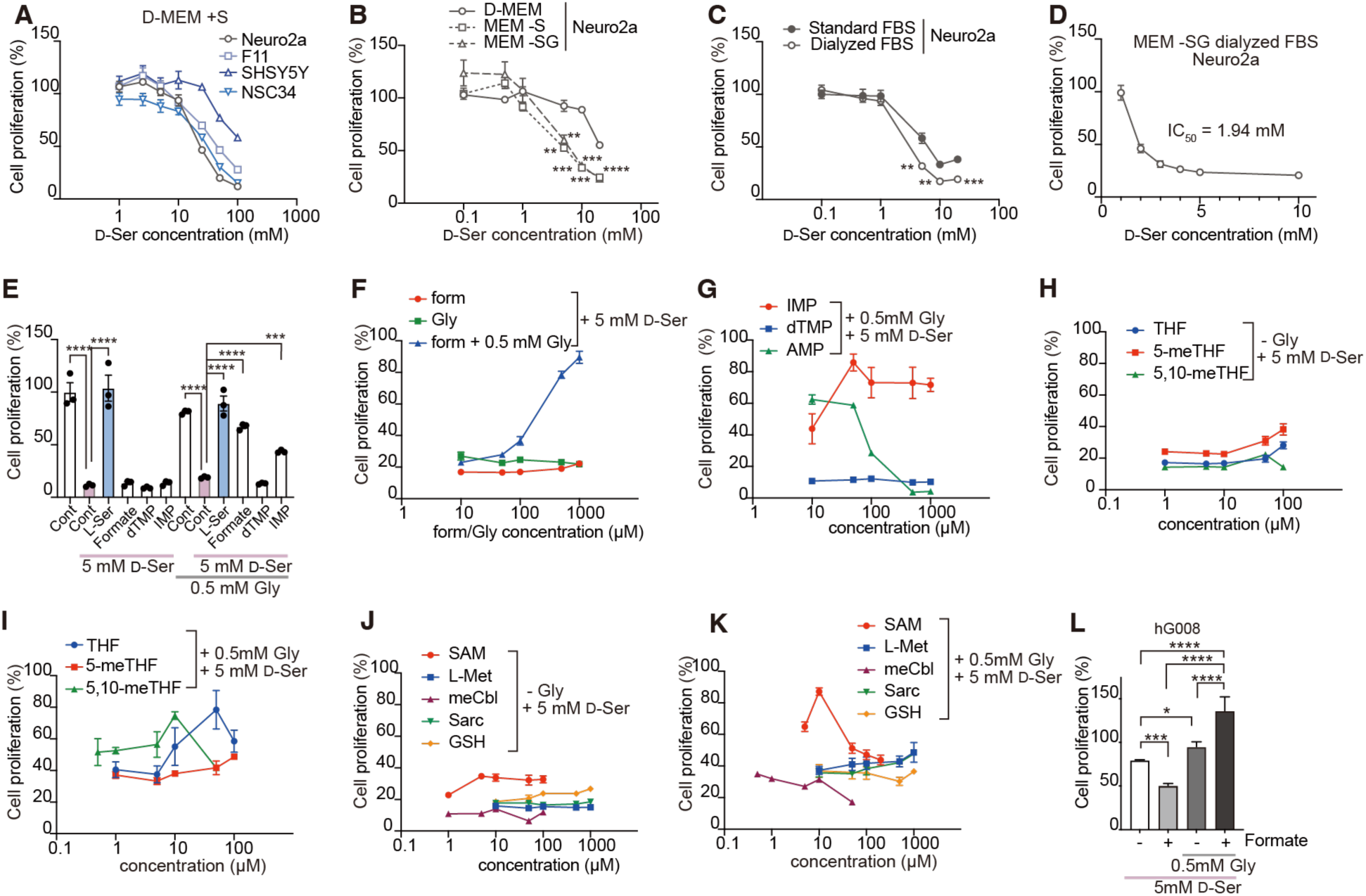
D-serine reduces tumor cell viability by inhibiting the folate cycle. **A.** Viabilities of neuroblastoma cell lines were cultured in D-MEM containing serine (+S) with the indicated concentration of D-serine (x-axis) (n = 4). **B.** Neuro2a Cells were cultured in D-MEM, MEM without serine (MEM -S), or MEM without serine/glycine (MEM -SG). **C.** Neuro2a Cells were cultured in MEM with 10% standard FBS or dialyzed FBS without serine/glycine. **D.** IC_50_ of D-serine on Neuro2a cell growth was analyzed in 10% dialyzed FBS, MEM -SG medium. **E-K.** Inhibitory effects of indicated metabolites against D-serine in the presence or absence of glycine were tested. Concentrations of amino acids or metabolites: D-serine (5 mM, **E-L**), glycine (0.5 mM, **E, G, I, K, L**), formate (1 mM, **E, L**), and dTMP/IMP (1 mM, **E**). **F** The indicated concentration of glycine (x-axis) was used for Gly group. The indicated concentration of formate (x-axis) was used for form group. 0.5 mM Gly and the indicated concentration of formate (x-axis) was used for form + Gly group. **L.** hG008 cells were cultured with the indicated concentration of D-serine and glycine (n = 4). Data are plotted as the mean ± s.e.m. Statistical significance was analyzed by one-way ANOVA. \**p* < 0.05, \*\**p* < 0.01, and \*\*\**p* < 0.001.

**Figure 4 figure supplement 6.**
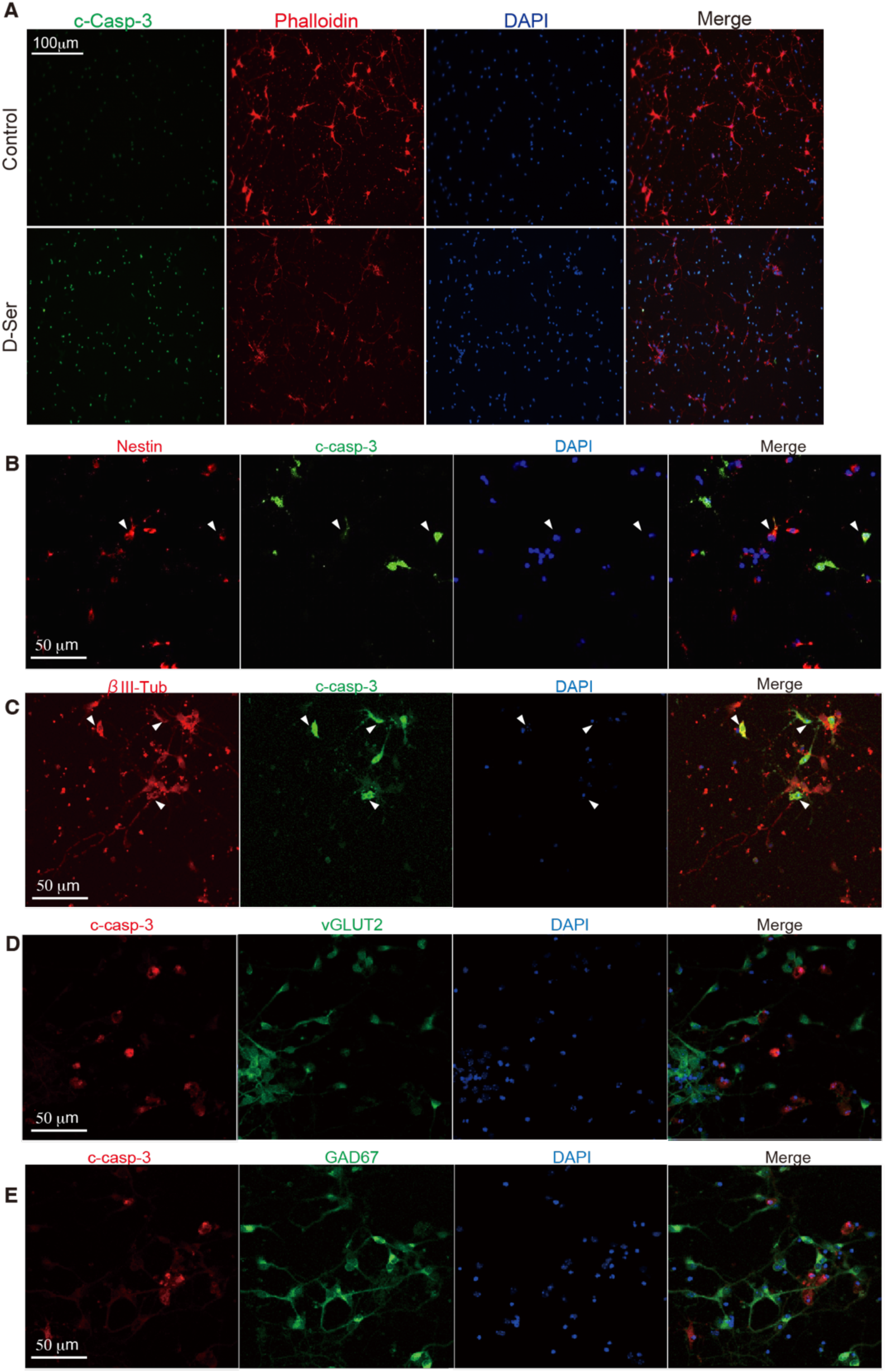
Immunofluorescence images for cleaved caspase-3 in NPCs treated with D-serine. **A** NPCs treated with control or D-serine for 48 hours were strained with DAPI (blue), an antibody against cleaved caspase-3 (c-Casp-3, green) and phalloidin (red). **B and C.** NPCs treated with D-serine for 48 hours were strained with DAPI (blue), antibodies against c-Casp-3 (green), and NPCs marker Nestin (red, **B**) or early neural marker beta III tubulin (βIII-Tub, red, **C**). Arrowheads show co-localization of c-Casp-3 and Nestin (B) or βIII-Tub (C). **D and E .** NPCs treated with D-serine for 48 hours were strained with DAPI (blue), antibodies against c-Casp-3 (red), and mature glutamatergic excitatory marker vGLUT2 (green, **D**) or mature GABAergic inhibitory marker GAD67 (green, **E**).

**Figure 4 figure supplement 7.**
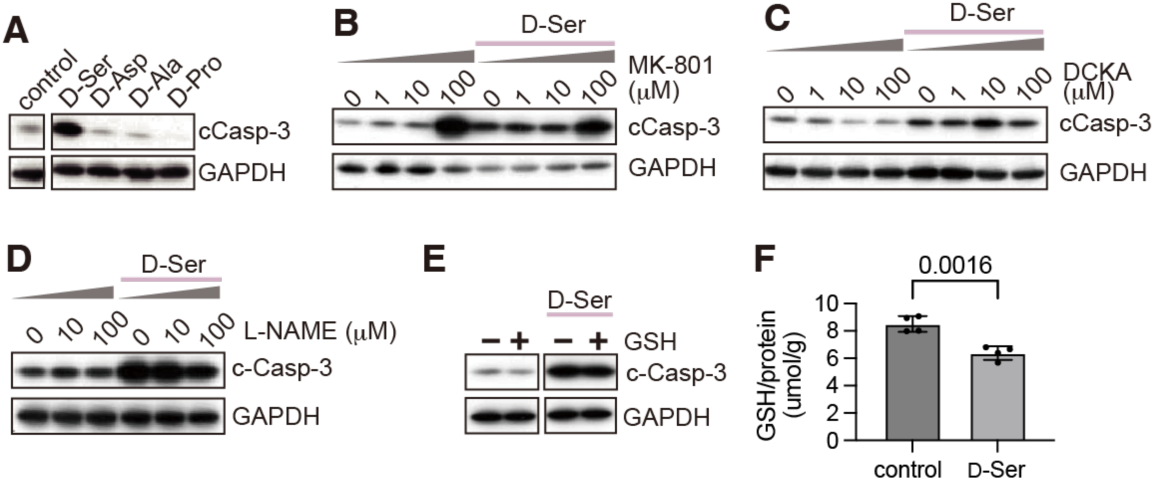
Characterization of apoptosis triggered by D-serine. **A.** NPCs were treated with D-amino acid for 48 h. **B-E**. Caspase-3 activation of NPCs treated with the indicated concentration of MK-801 (**B**) DCKA (**C**), L-NAME (**D**), or (**E**) with D-serine for 48 h. For (**B-E**), NPCs were treated with MK-801 (**B**), DCKA (**C**), L-NAME (**D**), or 100 μM glutathione (GSH) (**E**), 4 hours prior to D-serine treatment. Cleavage of caspase-3 (cCasp-3) in NPCs was detected using western blot analysis. (**F**) GSH concentrations in NPCs treated with D-serine for 48 hours were measured by enzymatic assay (n = 4). The concentration was normalized to total protein concentration.

**Figure 4 figure supplement 8.**
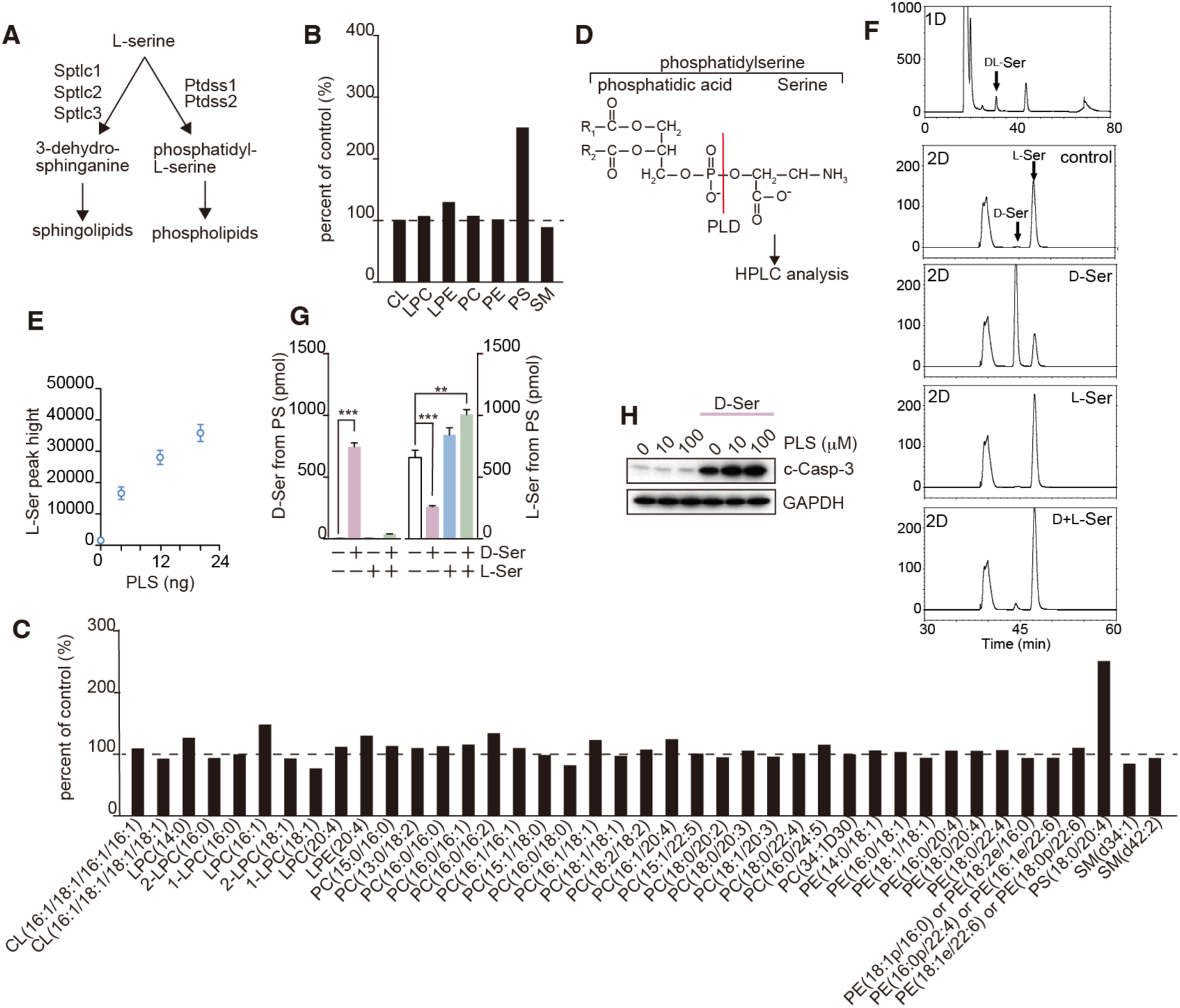
D-serine is incorporated into phosphatidylserine. **A**. A biosynthesis pathway for phospholipids and sphingolipids. L-serine serves as a component of both sphingolipids and phospholipids. **B**, **C**. Relative abundances of total cardiolipin (CL), lysophosphatidylcholine (LPC), phosphatidylcholine (PC), phosphatidylethanolamine (PE), phosphatidylserine (PS), and sphingomyelin (SM) (**B**) or single lipid molecules for each lipids (**C**) at 48h after treatment with D-serine, compared to untreated controls are presented (each group represents a pooled sample of n = 3). **D**. A cleavage site in PS by phospholipase D (PLD) is illustrated. D- or L-serine yielded by the cleavage was quantified using 2D HPLC. **E**. A standard curve for L-serine cleaved from phosphatidyl-L-serine (PLS) is shown. **F,G**. NPCs were treated with D- and/or L-serine. PS was extracted from NPCs and cleaved into D- or L-serine by PLD. Chromatograms show the first dimension (1D) and second dimension (2D) with loading time (min) and the signal intensity as X- and Y-axis, respectively (**F**). The peaks of DL-Ser in 1D, and D-Ser and L-Ser in 2D are indicated with arrows. Data with biological replicates (n = 4) are plotted (**G**) as the mean ± s.e.m. Statistical significance was analyzed by one-way ANOVA. **H**. Cleavage of Capsase-3 in NPCs treated with D-serine in the presence or absence of PLS at indicated doses were analyzed using western blot.

**Figure 4 figure supplement 9.**
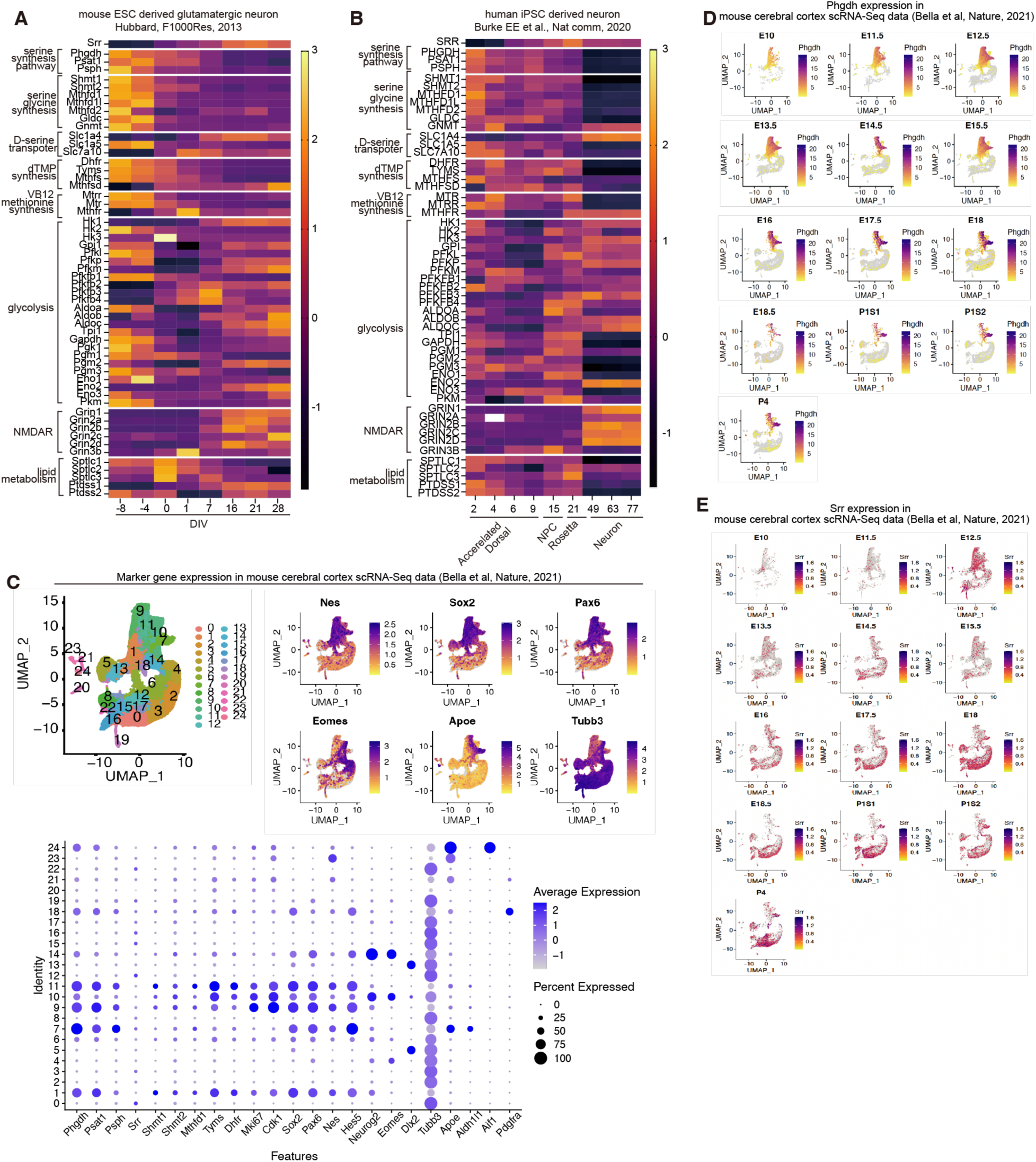
Transcriptional profiles of genes for serine metabolism during the development A, **B**. Genes related to L-serine metabolism were assembled and their transcriptional profiles along with development were analyzed using bulk RNA sequencing data from ES-derived (Hubbard et al., 2013) (**A**) or iPSC-derived neurons (Burke et al., 2020) (**B**). **C**. RNA expression levels of enzymes involved in the serine synthesis pathway, one-carbon metabolism, and folate cycle, and those of cell type markers for neural stem cells (Sox2), intermediate neural progenitors (Eomes), astrocytes (Apoe), and differentiated neurons (Tubb3) are shown as UMAPs and dot plots using single-cell transcriptome data from developing mouse brain by Bella et al(Bella et al., 2021). **D, E**. Developmental changes of expressional distributions of Phgdh (**D**) and Srr (**E**) are presented as UMAPs using data for the single-cell transcriptome.

**Figure 4 figure supplement 10.**
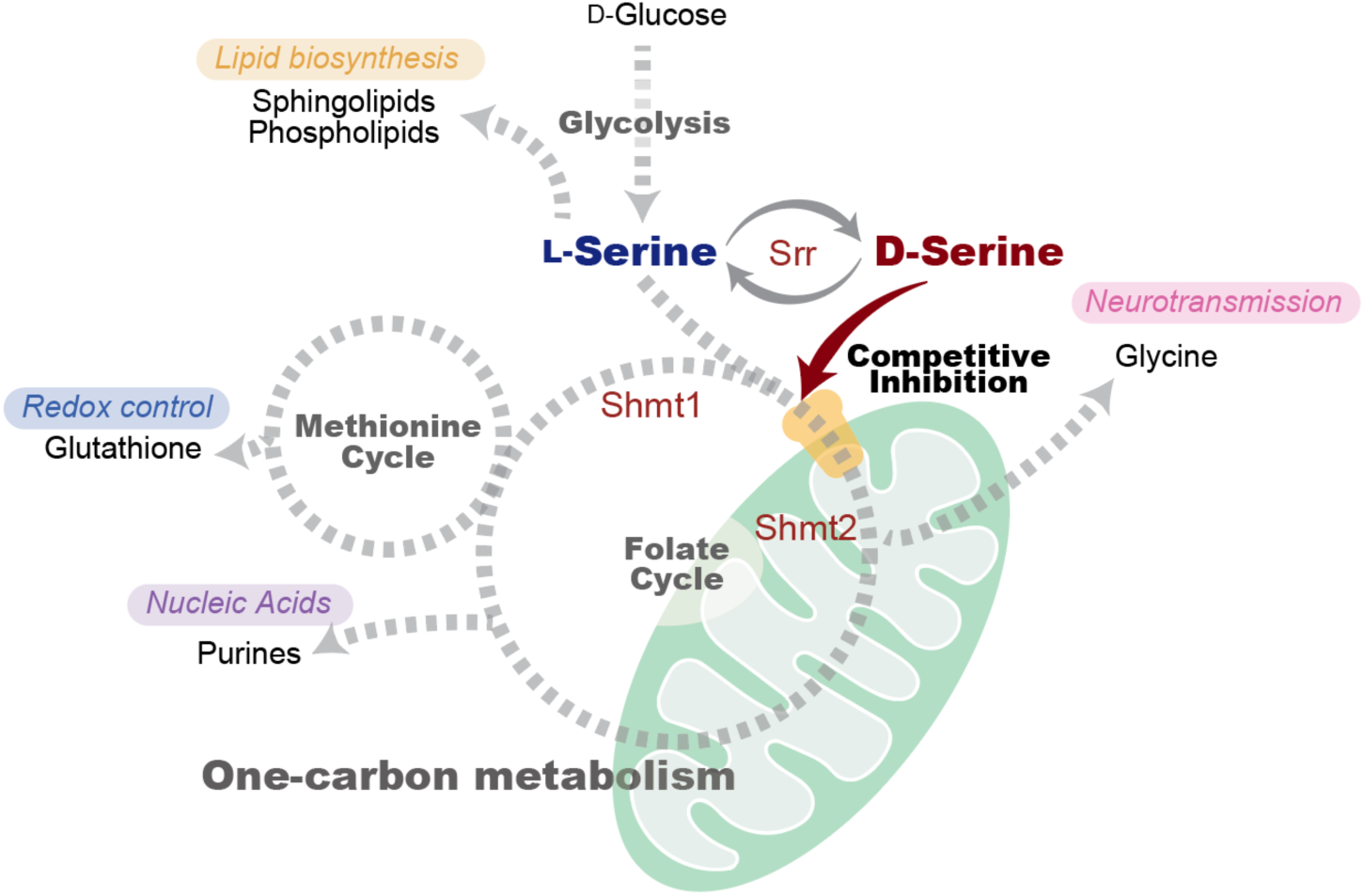
Schematic overview of the metabolic regulation by serine chirality. L-serine serves as a metabolic hub connecting glycolysis, lipid metabolism, and one-carbon metabolism, and is stereo-converted to D-serine by serine racemase (Srr) in the central nervous system. D-serine competes with L-serine for mitochondrial transport, thereby limiting mitochondrial L-serine availability and attenuating one-carbon metabolic flux. Under L-serine-limited conditions, this chiral competition restricts one-carbon metabolism-dependent cell proliferation.

